# Explanation of saltatory conduction in myelinated axons and of micro-saltatory conduction in C fibers of pain sensation by a wave-type ion plasmonic mechanism of stimulus kinetics

**DOI:** 10.1101/2023.07.14.548969

**Authors:** J. E. Jacak, W. A. Jacak

## Abstract

Ion plasmon-polariton model of stimulus in myelinated axons and in C fibers of pain sensation is developed. This solves a long standing problem in neuroscience of by 2 − 3 orders of magnitude discrepancy between the observed fast speed of the saltatory conduction in myelinated axons or in C fibers with the upper limit of diffusive ion current velocity in these axons. The latter, described in the framework of so-called cable model, is too low in axons because of poor conductivity of neuron inner cytosol. The compliance with observations has been achieved upon plasmonic model of ionic local oscillations synchronized in periodically corrugated axons and propagating with high speed in the form of wave-type plasmon-polariton without any net diffusion current, thus not limited by resistivity. The new model of stimulus in myelinated axons reveals the different controlling role of myelin than previously thought from cable model. The control mechanism in non-myelinated C fibers is also proposed in agreement with observations. Recognition of plasmon model of neural signaling may be important for identifying a new targets for the future treatment at demyelination diseases and for fighting pain.

## I. INTRODUCTION

Sufficiently fast signaling in neurons is of central significance for life functions of the body and for the functioning of the senses. Especially important is the high speed signal transduction in long myelinated axons in peripheral nervous system, which exhibits the velocity of action potential (AP) transduction of 100 − 150 m/s (in humans). Such rapid signaling is required to ensure a sufficiently quick reaction of the central nervous system needed to control distant organs and respond to signals from the senses conditioning survival in the environment. A lowering of the signal speed even by only 10 % ceases life functions. Myelinated axons prevail also in white matter of the brain and axon cord and such axons are typically longer than non-myelinated axons or dendrites of gray matter and also exhibit very fast signaling. AP is the carrier of information, which is able to ignite synapses in the whole electro-chemical paths in neural system. The transduction of AP along axons takes place via its repeating creation on trans-membrane ion channel complexes of Na^+^ and K^+^ according to well recognized Hodgkin-Huxley (HH) mechanism [1, 2], in the case of myelinated axons clustered at nodes of Ranvier – the breaks of ca 1 − 2 *μ*m length in myelin periodically distributed along axons in span of order of 100 *μ*m. HH cycle takes a time of order of 1 ms for AP creation and during this time period the signal with observed velocity of 100 m/s traverses 10 cm of axon length igniting ca. 1000 HH cycles on consecutive nodes of Ranvier. To ignite so large number of HH cycles and to shift AP along axon on the distance of 10 cm, the sufficiently fast stimulus is required. Thus the observed velocity of AP transduction is in fact the speed of the stimulus, which is the voltage signal with amplitude exceeding some threshold (in humans of 15 mV) causing local depolarization of the neuron membrane (from − 70 mV to − 55 mV) and triggering the opening of Na^+^ channels on consecutive nodes of Ranvier, which ignites HH cycles on these nodes. Conventionally, the diffusion current of ions along axons was considered as the stimulus signal, described upon the cable model in analogy to submarine telecommunication cable [3], as an axon is a tube with inner aqueous electrolyte (with molarity of order of 100 mM) isolated from similar inter-cellular electrolyte by insulating neuron membrane additionally covered by the lipid myelin sheath in the case of myelinated axons. The cable model well describes the stimulus kinetics in short naked dendrites [4, 5] or in non-myelinated axons [2, 6–11], where the Na^+^ and K^+^ channels are randomly distributed and not clustered as in myelinated axons at nodes of Ranvier. The speed of stimulus in dendrites or nonmyelinated axons is small, lower than 5 cm/s, as can be verified in the cable model for realistic electro-chemical parameters of neurons and the size and geometry of dendrites and nonmyelinated axons. The high speed of stimulus signal in myelinated axons is thus by three orders of magnitude larger than the ion diffusion velocity at the same axon diameter and even if increased ca. 10 times by the thick myelin (which lowers the electrostatic capacity *C* between inner and outer electrolyte across the membrane covered by myelin, and actually enhances the diffusion speed 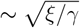 (*ξ* is the myelin layer thickness, *γ* is the naked membrane thickness) in cable model) is still by two orders larger and is called as the saltatory conduction without knowing a mechanism of so rapid jumping of the voltage signal between consecutive nodes of Ranvier [1, 9–19]. This is not a marginal problem in neuroscience, because the perturbation of signaling in myelinated axons is the source of severe diseases like Multiple Sclerosis or other demyelination syndromes and the absence of the recognition of physical mechanism of these malfunctions hampers a progress in related treatment. A loophole in understanding of saltatory conduction is also a serious problem in fundamental insight into functioning of nervous system, taking into account that majority of axons in brain and spinal cord are myelinated and form there the white matter and its role versus nonmyelinated filaments in gray matter is still puzzling.

The similar speedy communication as the saltatory conduction is also observed in naked nonmyelinated ultra-thin long axons of pain sensation – in so called C fibers for transmitting nociceptive pain sensations in peripheral neural system (cf. references in [20]), in which the AP transduction achieves the speed of 10 m/s, by three order of magnitude faster than the diffusion ion current in so thin (with diameter of order of 0.1 *μ*m) axons for which the cable model gives only 1.5 cm/s, additionally not increased by a myelin, which is absent here. The observed speedy signaling in C fibers is called as the micro-saltatory conduction without any imagination on its mechanism. The recent observations [20] revealed the periodic structure of sodium ion channels distribution along C fibers, where these channels are clustered on small thin lipid rafts of 0.1 − 0.3 *μ*m size and span periodically by 10 *μ*m. This structure resembles the organization of nodes of Ranvier in myelinated axons, though K^+^ channels are still randomly distributed along C fibers. Taking into account that Na^+^ channels are crucial for the ignition of HH mechanism of AP creation, one can infer that the periodic distribution of Na^+^ channel clusters is the fundamental property for speedy saltatory conduction both in C fibers and in myelinated axons, and the stimulus of such speedy firing of axons is not of diffusion current upon the cable model, but has a wave-type character.

In the present paper we develop a new model of fast stimulus kinetics in periodically modulated axons taking advantage of soft-plasmonics [21] related to synchronized ion density oscillations in aqueous electrolytes confined to micrometer size-scale by dielectric membranes, just as in the case of bio-cells. For elongate axons with periodic structure of myelin sheath or with periodicity of lipid rafts in C fibers, the wave-type plasmon-polariton excitation can be organized in the form of speedy propagating wave of synchronized ion oscillations (plasmons) in consecutive periodic segments. A wave packet of plasmon-polaritons in periodically corrugated axons can play the role of stimulus with the velocity highly exceeding the speed of diffusion of ions. Plasmon-polariton does not carry any net electrical current of ions and competes as stimulus with diffusive current of ions not perturbing the slower diffusion. However, only the fastest signal takes the role of a stimulus igniting HH cycles at consecutive periodic nodes. Because of the refractory period in HH cycles (i.e., a period of hundreds of milliseconds of node inactivity, which is needed to restore their steady state) only the fastest signal can ignite HH cycles, whereas slower ones are ineffective and are damped over short distances due to Ohmic losses without any stimulus significance, as they are not aided in energy on consecutive nodes. Only the fastest signal is supplemented in energy residually at HH cycles (on the eventual cost of ADP/ATP energy channel), which assures arbitrary long range of stimulus kinetics with the same amplitude. Plasmon-polariton is high frequency plasmonic-type oscillation of MHz frequency. Such e-m frequency signal in axons has not been subject of former studies, but recently has been observed in myelinated axons [22]. Wave type signal with e-m wave component as of plasmon-polariton can cross even small breaks in axons, which is impossible for diffusion current, but was observed in the case of two separated myelinated axons in pure water [23]. Both these observations [22, 23] are direct confirmations of plasmon-polariton stimulus kinetics. Moreover, other properties of plasmon-polariton – its size and temperature dependence, the external e-m signature (possible to be recognized by EEG or EMG techniques), also agree with observations besides just as needed velocity in myelinated axons and in C fibers.

Plasmon-polariton model of stimulus reveals a different role of myelin than previously thought from cable model. The thickness of the myelin sheath is crucial for the precise tuning of plasmon-polariton frequency to the time scale of triggering the opening of Na^+^ trans-membrane channels. The recent molecular study of these channels (on bacteria model [24]) allowed for the estimation of this timing to the scale of 1 *μ*s agreeing with MHz frequency of plasmon-polariton determining its velocity, on the other hand. The group velocity of plasmon-polariton wave packet is the higher the larger is its frequency, but too high frequency prevents the Na^+^ channels from opening. The observed speed of the saltatory conduction is the result of the trade-off between speed and frequency of plasmon-polariton. The mechanism of control over this velocity by myelin thickness is deciphered and the thinning or damage of the myelin sheath cause signaling perturbation due to desynchronization with opening of Na^+^ channels, but not due to an increase of electrostatic capacity across meyelinated cell membrane. The latter reduces the speed of the ion diffusion upon the cable model, which is still by two order of magnitude slower even at optimal myelin thickness, hence cannot be related with malfunctions of saltatory conduction. In C fibers, which are not myelinated, another mechanism of control over the plasmon-polariton micro-saltatory conduction has been identified and related with thickness of the bare axon. This elucidates why C fibers must be so thin, which on the other hand reduces the diffusion speed of ions in C fibers to only ca. 1 cm/s, thus without any significance for stimulus with velocity of 10 m/s observed in these axons.

The paper is organized as follows. In the next paragraph we formulate the mathematical model of plasmon-polariton in periodically corrugated axons. Next, the role of plasmon-polariton as stimulus for HH cycles in myelinated axons is discussed taking into account the molecular structure of Na^+^ channels. The related kinetics of plasmon-polariton stimulus is examined for realistic electro-chemical parameters of neurons and compared with observations. The control mechanism over plasmon-polariton by myelin thickness is presented, which allows for the simulation of realistic saltatory conduction. The generalization of the model for micro-saltatory conduction in C fibers is next formulated with the description of the control mechanism without a myelin. The view on the cable model and its insufficiency to describe saltatory conduction is shifted to Appendix.

## II. THE DERIVATION OF THE MODEL OF WAVE-TYPE SALTATORY CONDUCTION IN MYELINATED AXONS

We show how to solve a long-standing problem in neuroscience caused by a large discrepancy of two orders of magnitude between the conventional cable model and the firing rate observations of myelinated axons [1, 9–19]. The speed of stimulus responsible fo so-called saltatory conduction in myelinated axons cannot be elucidated by diffusion ion current in axons because of low velocity of the latter for poor conductivity if inner cytosol of neurons (cf. Appendix). Additional ca. 10-times increase of the velocity of diffusive current (upon the discrete HH cable model including reduction of trans-membrane capacity by myelin) diminishes the discrepancy with observations form three to two orders of magnitude. In the case of C fibers of pain sensation, which exhibit [20] also very fast micro-saltatory conduction, the discrepancy of the cable model velocity with observations remains on the level of three orders of magnitude, because of absence of a myelin sheath for C fibers. Apparently, the cable model (any ot its version) is not able to describe the saltatory and micro-saltatory conduction and other mechanism is responsible for fast kinetics of stimulus independently of too slow diffusion ion current. The common feature of myelinated axons and of C fibers is the periodic linear structure of an axon in both cases, which conditions a wave-type electro-signaling employing ion plasmons and plasmon-polaritons [25]. Such ion excitations are analogous to electronic plasmons in metallic nanoparticles [25–27] or electronic plasmon-polaritons in metallic nano-chains [28–30]. Due to larger mass of ions (by factor ∼ 10^4−5^) than of an electron, and much lower concentration of ions in dilute aqueous electrolytes in comparison to collectivized electrons in metals, the frequency of ion plasmons are by many orders of magnitude lower than of electron plasmons in metals and the scale of confinement suitable to manifestation of pronounced plasmonic effects is shifted from the nanoscale for electrons in metals to micrometer scale for electrolytes [21, 25]. Additionally, at lower than optical e-m frequency, the microwave (GHz) technique widely uses so-called Goubau lines [31], which very effectively transfer an electrical signal in a wave form in a periodically isolated metallic wires with electron oscillations, just like periodically corrugated myelinated axons with ion oscillations (at MHz frequency). The specific property of Goubau microwave signal is the possibility of its jumping across breaks in wire, in an air without any losses over large distances (even of several dozen of cm length), intentionally applied in related microwave Goubau circuits, and completely inaccessible for ordinary diffusion electrical currents. Electronic plasmon-polaritons and Goubau lines can be thus considered as archetype of ionic plasmon-polaritons in axons.

Ion plasmon-polariton can serve as the stimulus signal in periodically myelinated axons and in periodically corrugated (by lipid rafts) C fibers. The physics of plasmons and plasmon-polariton is, however, different from that of conventional diffusion current (in particular of ion currents upon the cable model – cf. Appendix), and need to be independently explored [25].

### A. Ionic plasmon-polariton in periodically corrugated axon

Myelinated axons differ from non-myelinated ones in their structure imposed by the periodic distribution of the myelin sheath – as illustration in Fig. 1. The length of myelinated segments – denoted by *L*, depends on the diameter *s* of axons, conserving the aspect ratio of order of 0.01, i.e., for most frequent in humans myelinated axons with diameter *s* = 1 *μ*m the myelin section length is ca. 100 *μ*m; the thickness of the myelin is for such an axon of order of *ξ* ≃ 0.2 *μ*m, the length of a naked node of Ranvier separating consecutive myelinated segments is ca. *δ* ≃ 1 − 2 *μ*m, whereas the thickness of a bare neuron membrane *γ* ≃ 7 nm. Size parameters of myelinated axons can vary to some extent [18, 32, 33], though in human majority of myelinated axons in central nervous systems are thinner than 1 *μ*m, and in peripheral neural system are averagely of 1 − 2 *μ*m diameter.

**FIG. 1.**
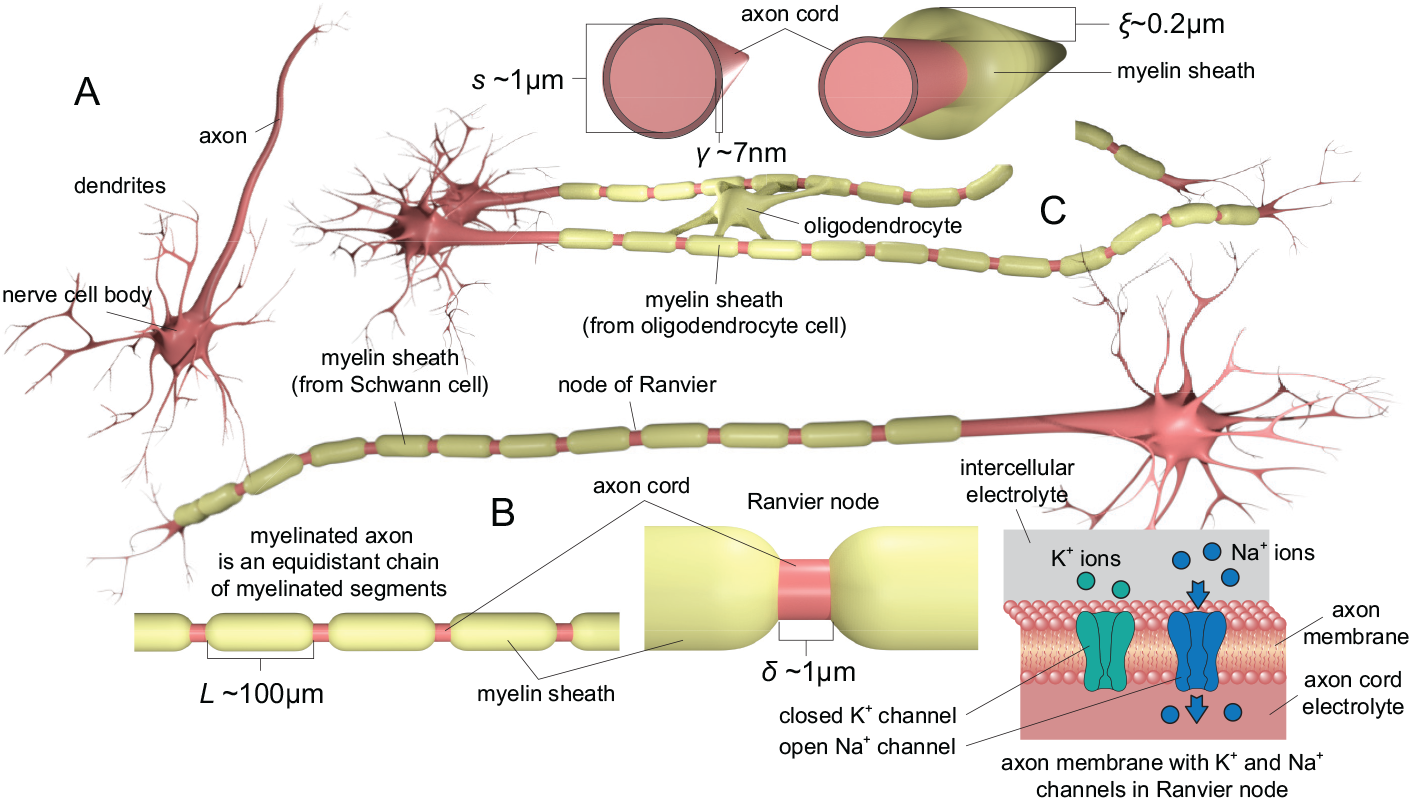
Illustration of axons: non-myelinated (A) and myelinated ones (B,C) – in white matter of the central nervous system myelinated by oligodendrocytes (C), in the peripheral nervous system by Schwann cells (B). Consecutive myelinated segments are separated by small breaks in the myelin – the so-called nodes of Ranvier. The length of the myelinated segment is marked by *L*, the diameter of inner cord by *s*, the myelin sheath thickness by *ξ*, the length of Ranvier node by *δ* (for example assumed *s* = 1 *μ*m, *ξ* = 0.2 *μ*m, *L* = 100 *μ*m, *δ* = 1 *μ*m, but these parameters can vary to a great extent). The naked membrane is thin – of thickness *γ* ≃ 7 nm. The myelinated axon resembles an equidistant chain of segments (lower left panel) – 10 cm of this chain contains ca. 1000 segments. The naked cellular membrane at nodes of Ranvier is pierced by trans-membrane ion Na^+^ and K^+^ channels gated by the membrane depolarization exceeding a certain threshold.

Let us consider the chain of myelinated axon segments separated by nodes of Ranvier – as shown in Figs 1 and 2.

**FIG. 2.**
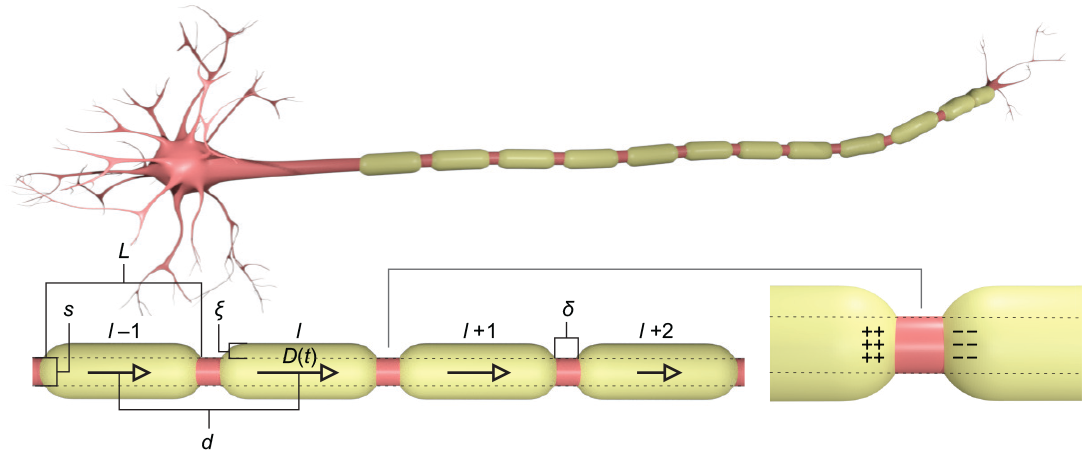
The myelinated axon can be considered as a chain of periodic segments, which can be numbered by consecutive integers …, *l* − 2, *l* − 1, *l, l* + 1, *l* + 2, … starting from an arbitrarily chosen *l*-th segment to the left and to the right (in the case of a finite axon the integers can change between some −*N* and *N* for *l* = 0 assigned to a segment in the middle of the axon). To each segment in its center can be pinned the local dipole of ion density oscillations *D*(*ld, t*) (*d* is the span between centers of neighboring segments), which varies along te chain in a wave-type according to Eq. (6). The dipoles in neighboring segments cause the time dependent longitudinal polarization of the node of Ranvier between these segments (lower panel), which induces the charge shifts in the node and the change of local depolarization of the membrane being the stimulus igniting the HH cycle.

In a single fragment of an xon cord below the myelin one can consider longitudinal dipole surface plasmon oscillations [25], i.e., synchronized local oscillations of ions (with concentration of ca. 100 mM in aqueous electrolyte of inner neuron cytosol). As was demonstrated in [21], ionic plasmons replicate electronic plasmons in metallic nano-particles, but in the scale of micrometer confinement agreeing with size of axon segments. The frequency of the longitudinal local dipole oscillations of ions we denote by *ω*_1_.

The oscillating dipole of local ion density in a particular segment, pinned to the center of the selected segment of an axon (denoted by the vector **r**, in arbitrarily chosen coordinate system), induces also ion oscillations in neighboring segments via electric coupling between oscillating charges. This coupling can be described by the electric field emitted by the oscillating dipole in point **r** according to the formula [34, 35] for the electric field induced in point **r** + **r**_0_ (for arbitrary vector **r**_0_),

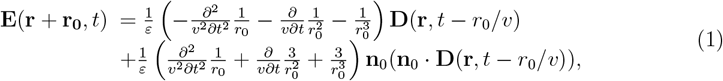

where 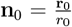 (*r*_0_ is the length of the vector **r**_0_). Note that the field **E** in the point **r** + **r**_0_ at time instant *t* had been induced earlier by the dipole **D** at **r** – at time instant *t* − *r*_0_*/v*, because of light velocity limit for any e-m signal propagation (here 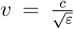 is the light velocity in the medium with the permittivity *ε* and *c* = 3 × 10^8^ m/s is the light speed in vacuum).

In Eq. (1) the terms with denominators of 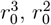, and *r*_0_ are referred to as the near-field, medium-field, and far-field zone components of the dipole radiation, respectively, [34, 35], because terms with the denominator 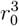 are largest at small distances from the point **r**, whereas terms with *r*_0_ denominator prevail at greater distances. Eq. (1) allows to quantitatively express the mutual interaction of oscillating ion dipoles on the segments in the axon – the dipole on each segment interacts with all other segments, most strongly with the closest neighbors and weaker with next-nearest neighbors, but the interaction of all must be included.

Because of linear alignment of axon segments one can choose **r** = 0, i.e., the coordinate origin can be associated with the center of a selected segment. Then, **r**_0_ vector to the center of an arbitrary other segment is parallel to the axon axis (assumed here in *z* direction) and can be assigned by the integer *l* and *r*_0_ = *ld*, where *d* is the distance between centers of consecutive segments (*l* can vary between − ∞ and +∞ for theoretically infinitely long axon).

The segments in the chain with own dipoles of ion local oscillations can be numbered by consecutive integers (cf. Fig. 2), and the equation for the dipole oscillations of the *l*-th segment can be written as follows,

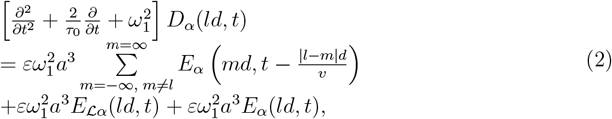

where *α* = *z* indicates the longitudinal polarization of dipole oscillations with respect to the axon orientation (assumed along *z* axis), whereas *α* = *x*(*y*) the transverse polarization (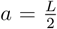, where *L* is the length of a myelinated segment, note that *d* = *L* + *δ*). Summation over integer *m* represents the interaction between dipoles located on all others segments in the chain with the dipole at selected *l*-th segment. If one omits the right-hand-side of Eq. (2), the simple harmonic oscillator equation for a dipole on *l*-segment remains,

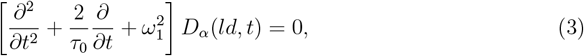

i.e., the harmonic oscillator equation for the dipole *D*_*α*_(*ld, t*) on the *l*-th segment of the axon with the self-frequency *ω*_1_ and with damping 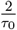 caused by ordinary Ohmic losses due to ion scattering at oscillations. The first term on the right-hand side of Eq. (2) is the dipole coupling due to electric field radiation from all other segments. The second term on the right-hand side of Eq. (2) describes the *l*-th dipole damping due to radiation losses. Such losses are caused by the outflux of energy of radiating oscillating charges, which can be represented by the inhibiting effective field *E*_ℒ_ acting on the dipole *D* in point *ld*. This effective field resembles a friction field dissipating energy of oscillations and is called as Lorentz friction, [34, 35]. The third term on the right-hand side of Eq. (2) represents the external forcing field inducing oscillations of ions in the chain.

We will focus our attention on the longitudinal dipole oscillations, because for the elongate geometry of axon segments such oscillating dipoles in consecutive segments under myelin cause an effective dipole oppositely oriented along the node of Ranvier (as shown in Fig. 2). The charge shifts at nodes of Ranvier resulting from longitudinal dipole ion oscillations in adjacent myelinated segments are responsible for the triggering of the HH cycles on these nodes.

Ohmic losses due to scattering of ions during oscillations in the single segment can be expressed via the term,

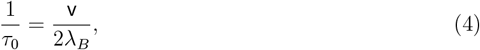

where *λ*_*B*_ is the mean free path of ions in the electrolyte (as in bulk), v is the mean velocity of ions at the temperature *T* (in Kelvin scale), 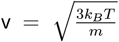, *m* is the mass of the ion (assuming one kind of ions for a model), *k*_*B*_ is the Boltzmann constant. Note, that this is a place where the temperature influences the plasmon-polariton kinetics, according to Eq. (4) higher temperature rises damping of ion oscillations 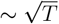.

Using Eq. (1) it is possible to express the first term on the right-hand size of Eq. (2) in the following form (for longitudinal polarization of ion oscillations i.e., for *α* = *z*),

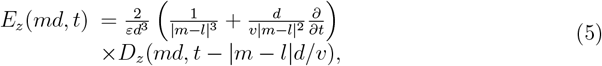

Plasmon-polariton is the solution of Eq. (2), which would display the collective ion oscillations in all segments in the chain, synchronized mutually via electrical coupling expressed by Eq. (5). Eq. (2) is a linear differential equation with respect to *D*_*z*_ both in the position and time domains. Position variable is discrete expressed by integers *l* ∈(− ∞, ∞), whereas time *t* is continuous. Moreover, the position variable displays periodicity of segments alignment in an axon. Standard method for the solution of such a differential equation is taking the Fourier picture in both domains, which converts the differential equation into algebraic one. Because of periodicity in the chain the position dependent Fourier transform acquires the form of Fourier series, or, in other words, must be a discrete Fourier transform with related Fourier counterpart of discrete position in the form of wave vector (in 1D one dimensional vector, i.e., scalar) 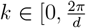), like in the case of 1D crystal with period *d*. Hence, the single Fourier component (discrete for position domain and continuous for time domain) must be of wave type form displaying the single mode of the collective synchronized oscillations in the whole axon propagating as a wave,

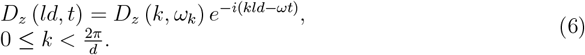

Note that *k* must be taken form the finite set [0, 2*π/d*) because of periodicity and for infinite chain is a continuous variable, though for a finite chain must be discrete 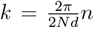, *n* = 0, …, 2*N* assuming here 2*N* + 1 segments in the axon with length ℒ = (2*N* + 1)*d*. The boundary condition (of Born-Karman-type) imposed on the entire system, *D*_*z*_(*ld* +ℒ) = *D*_*z*_(*ld*), results in the discrete possible values of *k*, i.e., 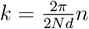, *n* = 0, …, 2*N*. If one takes the limit *N* → ∞ (infinite chain), the variable *k* becomes continuous, although *k* ∈ [0, 2*π/d*) still holds. The variable *ω* representing the time dependent Fourier transform is not restricted by any constraints (because of continuity of time). The wave type single mode solution (6) must be substituted to the equation Eq. (2) and represents a specific form of its solution because of the linearity of Eq. (2). The general solution is of the form of linear combination of single Fourier modes and is called as a wave packet of plasmon-polaritons. This is a common behavior of all oscillatory signals like visual signals or sounds, which are wave packets of photons or phonons (single Fourier components), respectively. For each admissible value of *k* wave vector (we use traditional name ‘wave vector’ despite it is a scalar in 1D case) the solution of Eq. (2) gives the related *ω*_*k*_. *k* numerates thus various modes of the wave-type solution called here plasmon-polaritons (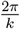 is the wavelength of the mode *k* of plasmon-polariton) (6), whereas corresponding *ω*_*k*_ defines the frequency, *Reω*_*k*_ of *k*-mode of plasmon-polariton and its damping *Imω*_*k*_ (because Eq. (2) has nonzero its imaginary part, due to the term with 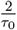 and also imaginary part of the right-hand side of the equation – thus *ω* must be a complex variable, at assumption that *k* is real).

The Fourier picture of Eq. (2) (for *α* = *z* – longitudinal plasmon-polariton) attains thus the form,

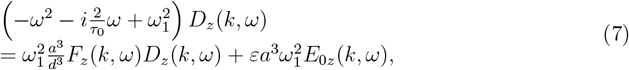

with

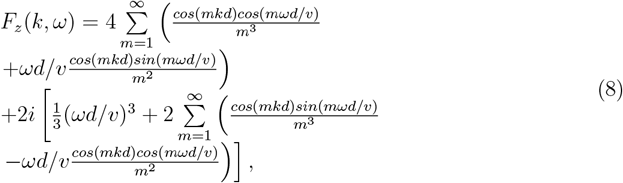

note that the Lorentz friction is imaginary (as damping term) and is included to the imaginary part of *F*_*z*_ as the first term in its imaginary part (because for an oscillating dipole 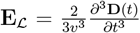 [34, 35]). The trigonometric functions in Eq. (8) arisen due to summation of antisymmetric Fourier-type terms for *m* and − *m*, which simultaneously reduced infinite sums to only positive integers *m*.

In order to find modes of plasmon-polariton in the myelinated axon, one must solve Eq. (7) with respect to *ω* for each value of *k* ∈ [0, 2*π/d*) (the variable *ω* is a complex number, in general). As mentioned above, the real part of the solution for *ω* will determine the frequency of plasmon-polariton mode *Reω*_*k*_, whereas the imaginary part *Imω*_*k*_ will describe the damping of this mode. Note that the function *Reω*_*k*_ (next we will call it simply *ω*_*k*_, referring to plasmon-polariton damping separately – cf. next paragraph) forms a band of modes (like in 1D crystal). The band of these frequencies looks like that presented in Fig. 3 (left panels) where the exemplary solutions of Eq. (7) are shown.

**FIG. 3.**
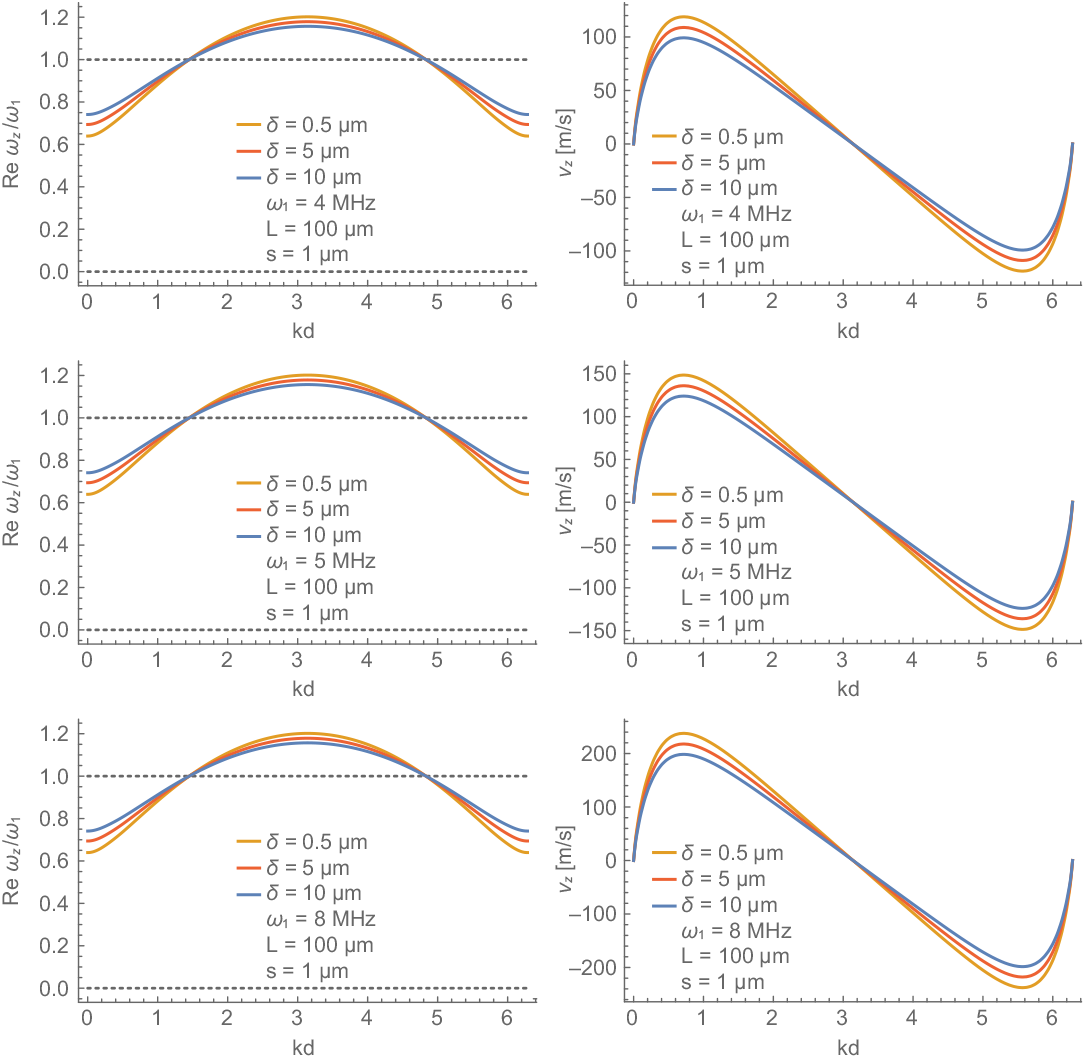
Solution of Eq. (7) for longitudinal plasmon-polariton mode frequency *ω*_*z*_(*k*) (in units of *ω*_1_ – the self-frequency of longitudinal surface plasmons in a single segment, if separated) with respect to wave number *k* (*kd* ∈ [0, 2*π*), *d* is the distance between centers of consecutive myelinated segments (*d* = *L* + *δ*), i.e., the periodicity scale of the structure), and the corresponding group velocity 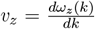 (in units of m/s) for axon parameters (for the length of myelinated segments *L* = 100 *μ*m, and the length of nodes of Ranvier *δ* = 0.5, 5, 10 *μ*m, for comparison). Three values of *ω*_1_ (the self-frequency of plasmons on a single segment, if separated) are assumed (for comparison), *ω*_1_ = 4, 5, 8 MHz. Strong dependence on *ω*_1_ is visible. The dependence on *δ* (the length of Ranvier node) is weak. For *ω*_1_ = 5 MHz (middle panel), the synchronization of the plasmon-polariton frequency with time-scale of 1.5 *μ*s for triggering of sodium channels opening takes place, which leads to the stabilization of this mode of plasmon-polariton with *v*_*z*_ agreeing to observed saltatory conduction speed for such axon parameters. It was assumed here the aspect ratio 0.01 of myelinated segment (i.e., the axon cord diameter *s* = 1 *μ*m). The myelin sheath thickness was assumed *ξ* = 0.2 *μ*m. The self-frequency *ω*_*z*_(*k*) is the real part of the solution of Eq. (7) for complex in general *ω*, i.e., *Reω*_*z*_(*k*) (normalized by *ω*_1_); 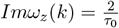 because of vanishing of radiation losses, cf. Eq. (9).

## III. PROPERTIES OF PLASMON-POLARITON STIMULUS KINETICS IN AXONS

### A. Absence of external e-m signature of plasmon-polariton

By the direct summation in Eq. (8) one can find that

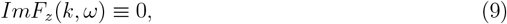

which means the absence of the radiation losses of plasmon-polariton in the axon (the imaginary part of *ω* will be thus determined by Ohmic losses only, i.e., by 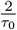 term). In other words, the amount of the energy incoming from other segments in the chain to a particular segment (via the dipole interaction in axon) is exactly the same as the energy outflow from this segment due to individual radiation (i.e., the Lorentz friction in all segments is fully balanced in periodically aligned segments of the axon). This means that ionic plasmon-polariton (similarly as electronic plasmon-polariton [25]) has no external e-m signature. The saltatory conduction is thus energy frugal – does not dissipate energy in a radiative way.

One can easy verify this result taking analytically infinite sums contributing to the imaginary part of *F*_*z*_ in Eq. (8), according to the following exact formulae [36],

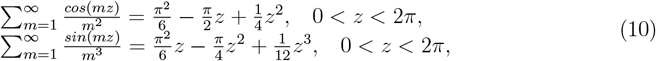

which gives (from Eqs (8), taking into account trigonometric relations, 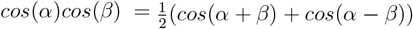 and 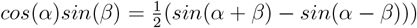 the Eq. (9).

The irreversible losses of energy of plasmon-polariton are linked only to scattering of ions (the term with 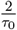), which give Ohmic losses eventually converted into Joule heat. Moreover, dipole oscillations in segments of an axon couple to inter-cellular electrolyte, which causes additional nonradiative losses and energy dissipation, small for myelinated axons because of thick myelin sheath, but large in C fibers, which are not isolated from surroundings by a myelin layer (this will be discussed further).

Though the result (9) is accurate for infinite axon (the sums are infinite), for a finite axons the property (9) is almost perfectly satisfied with high accuracy because of very quick convergence of series (10) contributing to the *ImF*_*z*_ – even only first 10 terms in these series give almost exact sums. Hence, the plasmon-polariton kinetics in short finite axons is very well approximated by its model in infinite chains.

### B. Frequency and speed of plasmon-polariton

Eq. (7), though linear differential equation with respect to *D*_*z*_, is highly nonlinear with respect to the complex *ω* and can be solved both perturbatively in an analytical manner [37] or numerically beyond the perturbation approach [38]. The solutions for *ω*_*k*_ can be applied to the myelinated axon model if the parameters in the dynamic equation (7) are adjusted to the actual neuron parameters.

The speed of the plasmon-polariton mode is given by the derivative 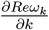, which is called the group velocity *v*_*g*_(*k*) of plasmon-polariton signal (cf. paragraph III C for explanation of the group velocity of a wave packet). The group velocity depends on the wave vector (here number, for 1D case) *k* numerating modes. From single modes it can be formed the wave packet of plasmon-polaritons, which propagates with the velocity equal to the group velocity of the central component of the packet. The exemplary solution of Eq. (7) are plotted in Fig. 3 for *Reω*_*k*_ (for short, denoted as *ω*_*k*_) with corresponding group velocities. Damping of the mode *k* of plasmon-polariton is equal to 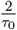 (cf. Eq. (4)), i.e., to Ohmic losses damping rate, because of the vanishing of radiation losses (additional damping due to coupling with intercellular electrolyte in the case of C fibers will be commented further).

The presented in this paper simulation (cf. Table I) of the proposed mechanism of myelinated axon firing by the ignition of the HH cycles on consecutive nodes of Ranvier by the wave-type plasmon-polariton stimulus signal instead of the conventional diffusive current, evidences the signal transduction as speedy as that observed at the saltatory conduction. The model is non-local and concerns synchronized ionic oscillations in periodically distributed segments of axons, which can propagate as a wave packet without a net current along the arbitrarily long axon with high speed provided that the frequency of ion oscillations is synchronized with the timing of the triggering of sodium channels at nodes of Ranvier, i.e., is of order of 4 MHz (this particular value of plasmon-polariton frequency will be clarified in paragraph IV B). The supporting argument is a recent observation of just such frequency of e-m oscillations in myelinated axons [22]. Another, relatively old observation of the jump of the ignition signal between separated (in water) two axons [23], also suggests the existence of wave-type component of signaling in myelinated axons as in the proposed plasmon-polariton model (any jump of signal across a gap filled with water is precluded for ion diffusive currents within the cable model approach).

**TABLE I.** Simulation within the plasmon-polariton approach (numerical solution of Eqs (7) and at plasmon-polariton frequency *ω* synchronized with HH cycle^*\**^, simulation for 10^4^ nodes)

| $s$ [$\mu\text{m}$] | $\xi$ [$\mu\text{m}$] | $L$ [$\mu\text{m}$] | $\delta$ [$\mu\text{m}$] | $\omega_1$ [MHz] | $T$ [$\mu\text{s}$] | $v$ [m/s] |
| --- | --- | --- | --- | --- | --- | --- |
| 1 | 0.23 | 100 | 2 | 5 | 1.5 | 140 |
| 0.8 | 0.2 | 100 | 1 | 4.2 | 1.9 | 110 |
| 1 | 0.2 | 120 | 2 | 5 | 1.5 | 160 |
| 1 | 0.2 | 100 | 2 | 3.5 | 2.3 | 100 |
| 1 | 0.2 | 100 | 1 | 3.1 | 2.6 | 90 |
| 1 | 0.2 | 100 | 1 | 2.1 | 3.8 | 60 |
| 1 | 0.2 | 120 | 1.2 | 2.1 | 3.8 | 75 |
| 1 | 0.23 | 100 | 1 | 0.62 | 12.5 | 19 |
| 1 | 0.23 | 180 | 1.5 | 0.62 | 12.5 | 32 |
| 1 | 0.2 | 100 | 2 | 8 | 1 | 200 |

\* $\omega = \frac{2\pi}{T}$ , ( $\omega$ is the solution of Eq. (7) for assumed $\omega_1$ and taken at $k$ for maximal group velocity $\frac{d\omega(k)}{dk}$ ), $T$ gives the corresponding needed lowest time scale for triggering of $\text{Na}^+$ channels at the HH cycle ignition;
list of symbols: $s$ – inner axon diameter, $\xi$ – myelin layer thickness, $L$ – internodal length, $\delta$ – node length, $\omega_1$ – frequency of plasmon oscillations in a single segment when is separated, controlled by $\xi$ , $T$ – time-scale of triggering of $\text{Na}^+$ channels required for plasmon-polariton at assumed $\omega_1$ , $v$ – speed of plasmon-polariton wave packet;
the velocity of plasmon-polariton signal scales with the axon diameter linearly, $\sim s$ .

### C. The wave packet of plasmon-polaritons and its group velocity

Plasmon-polaritons defined by single Fourier components (6) are specific solutions of the wave-type equation (7), when forcing external field is zero. Such a homogeneous differential wave equation defines the self-modes of waves, called here plasmon-polaritons. Due to linearity of this equation with respect to *D*_*z*_, any linear combination of self-modes is also the solution. Fourier-form self-modes are not localized in time and space, i.e., their oscillations span uniformly over the whole time and space domains. The sums of self-modes (integrals over some subdomain of wave vectors) form so-called wave-packets, which are localized in time and space and display the movement of plasmon-polariton signals in a similar manner as signals of light or sound being also wave packets of not localized self-modes for photons or phonons.

To be more specific, the self-modes including those given by Eq. (6), depend on time and position via the exponent 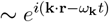, where **k** is the wave vector (3D in general case) numerating modes and *ω*_**k**_ is the frequency of **k** mode. In 1D case, as for plasmon-polaritons in axons, the wave vector is simply a number and position is discrete. The phase of such a single mode, **k** · **r** − *ω*_**k**_*t*, defines the variation of the signal in space and time, though the oscillations of this mode span uniformly in the whole position space and for *t* ∈ (− ∞, ∞). For single self-mode the phase velocity can be defined as the speed of an external observer accompanying the constant phase of the exponent, **k** · **r** − *ω*_**k**_*t* = *const*., from which (after taking the derivative with respect to *t*), 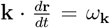. The velocity of the observer, 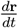, in direction of **k** is called as the phase velocity and equals to 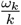 where *k* = |**k**|.

The general solution of wave-equation is the linear combination of single Fourier modes – the integer over some subset of self-modes with the envelope function, which selects particular single modes to the sum. The resulting signal is usually localized in time and space (it displays a moving hillock of the amplitude) and is suitable to transfer signals (energy and information) – is called as a wave packet and attains the following form (for simplicity written here for 1D case of wave-type kinetic along *z* direction, like in axons),

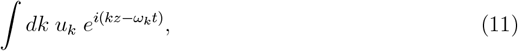

where *u*_*k*_ is the envelope function defining weights for particular modes *k* contributing to the wave packet. If *u*_*k*_ is nonzero on some region Δ*k* and concentrated (maximal) around certain *k*_0_, then the wave packet (11) can be rewritten as follows (with accuracy to linear departure from *k*_0_),

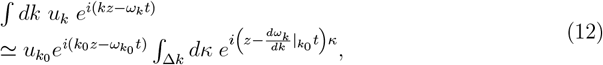

where *κ* = *k* ℒ *k*_0_.

The last integral in (12) defines the resulting amplitude of the wave packet, which vanishes (is very small) everywhere with exception to 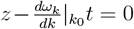, i.e., this is nonzero hillock of the amplitude, which moves along *z* direction with the velocity 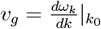. This velocity is called the group velocity of the wave packet. Usually it is the group velocity of the central mode of the wave packet. The group velocity is different than the phase velocity, unless the frequency spectrum is linear (as for photons). For plasmon-polaritons the frequency spectrum in the band is nonlinear, thus the group velocity is different than the phase velocity of a particular mode. The envelope function is shaped by the external forcing field contributing to the wave equation by unhomogeneous term.

Transfer of signals by wave packets is a common behavior of all wave-type phenomena including visual or sound signals localized in space and time and built of non-localized plane waves of photons or phonons and can be applied to wave packet of plasmon-polaritons being the carrier of stimulus signal in axons.

The group velocity 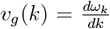 can be associated to any wave vector *k* ∈ [0, 2*π/d*) and can represent the group velocity of a wave packet when a particular *k* is a central point of an envelope function appropriately selected by external factors.

## IV. CONTROL OVER PLASMON-POLARITON IN PERIODICALLY CORRUGATED AXONS

### A. Coupling of plasmon-polariton with HH cycles

In any mechanism of AP signal transduction along axons, the most important role play HH cycles [1, 2] at nodes of Ranvier (in myelinated axons) or at small lipid rafts with Na^+^ channels (in C fibers) [20]. HH mechanism ignited by a stimulus signal reproduces AP at each of consecutive nodes with time retardation resulted from the finite speed of stimulus. Hence, the AP – the carrier of information in neurons, traverses along axon with the velocity of the stimulus signal. The AP with precisely repeatable shape is creating due to the activity of trans-membrane channels for Na^+^ (and K^+^) ions ignited by a stimulus voltage signal exceeding some threshold value. At steady conditions the concentration of ions Na^+^ is 10 times larger outside than inside of an axon and of ions K^+^, inversely. The polarization of an axon membrane at the steady state is (in humans) about −70 mV. An increase of this voltage to −55 mV by the stimulus signal opens Na^+^ channels (the molecular mechanism of this opening will be detailed in paragraph IV B) and causes an influx of Na^+^ ions to inside of the axon, which increases the membrane depolarization to about +30 mV. At such a voltage across the membrane the opening of K^+^ channels takes place, which stops further depolarization and allows rapid outflux of K^+^ ions outside the axon along the gradient of their concentration. This phase called as repolarization reduces the trans-membrane voltage again to −70 mV, with even some hyperpolarization by additional 10 − 15 mV, which, however, gradually disappears in next few milliseconds and the steady state is restored. The recovery of a steady state needs, however, an active transport of Na^+^ and K^+^ ions across the membrane against the concentration gradient. Two phases, the depolarization and the repolarization take together ca. 1 ms, whereas the stimulus action which triggers the opening of sodium channels is much shorter – sub-millisecond [24, 39–41]. The restoration of the steady conditions from the hyperpolarized state is energy-cost and takes place via the ATP/ADP mechanism (phases of HH cycle are shown in Fig. 5). Each ATP-phase cycle transports one positive elementary charge contrary to Na^+^ and K^+^ concentration gradients across the axon membrane. This process saturates for a longer time (even up to 1 s). Thus an expired block of Na^+^ and K^+^ ion channels (e.g., at a node of Ranvier) is inactive (called as refractory period) up to the restoration of its steady state.

**FIG. 4.**
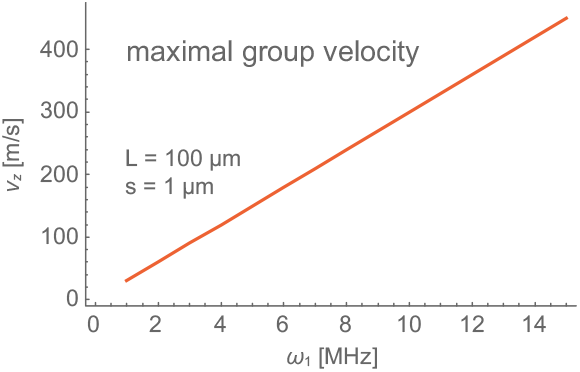
Dependence of the maximal group velocity *v*_*z*_ (in m/s units) of longitudinal plasmon-polariton mode with respect to the self-frequency *ω*_1_ (in MHz units) of plasmons in single myelinated segment (the segment length *L* = 100 *μ*m, the length of Ranvier node *δ* = 0.5 *μ*m, the diameter of the axon cord *s* = 1 *μ*m, the thickness of the myelin sheath *ξ* = 0.2 *μ*m).

**FIG. 5.**
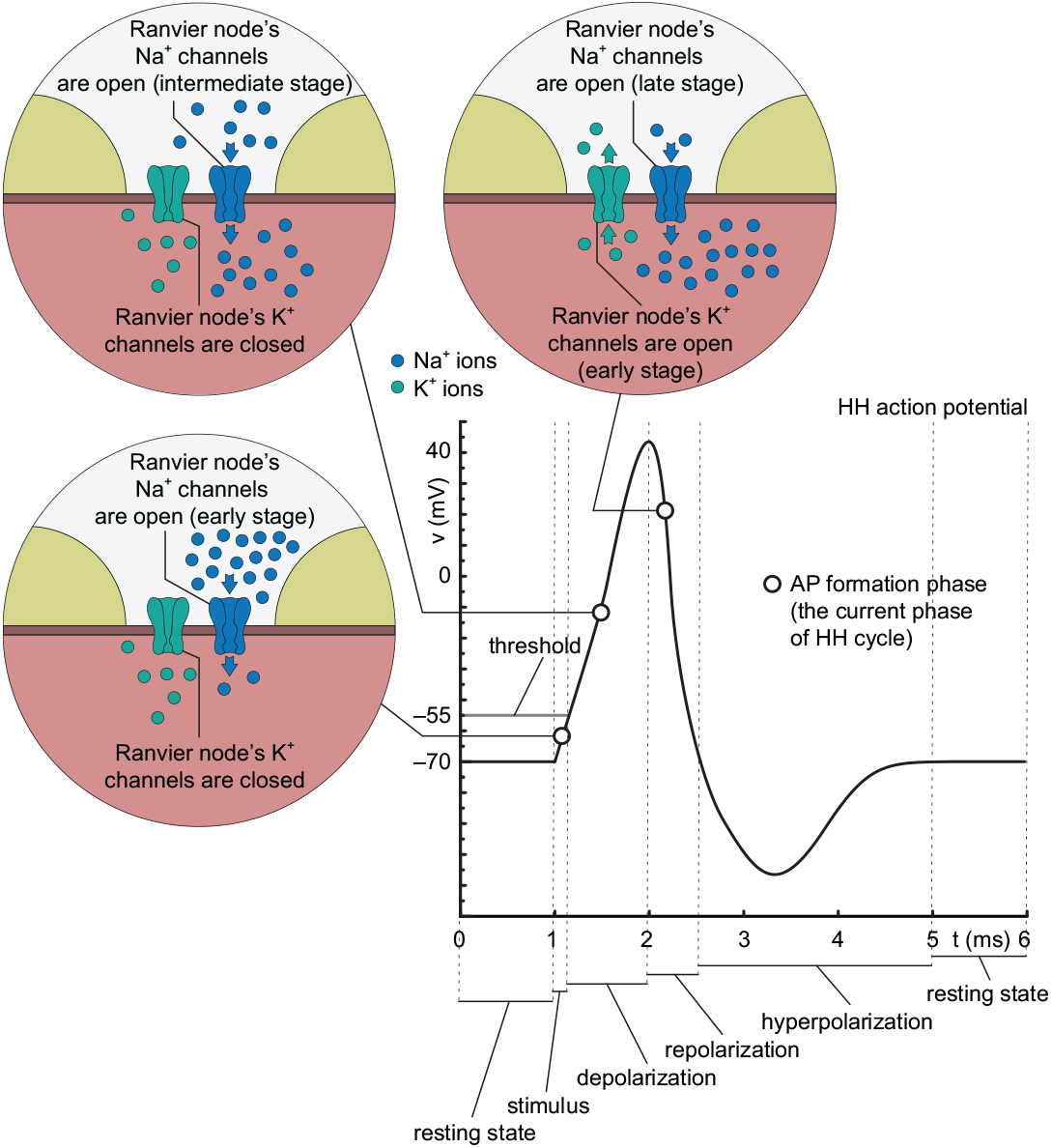
AP formation begins with the opening of trans-membrane sodium channels crossing the naked cell membrane at the nodes of Ranvier. The opening of sodium channels is triggered by the stimulus signal, which propagates with finite speed (in the case of myelinated axons, with the speed of 100 − 150 m/s). Hence, the formation of the AP on the consecutive nodes of Ranvier is in various phases of the HH cycle at distinct nodes – the time retardation is caused by the finite speed of the stimulus signal. In the upper panel the initial step is visualized – the Na^+^ channels are just opened and an influx of Na^+^ ions from the outside to the inner part of the axon starts, the K^+^ channels are still closed. Middle panel shows an intermediate phase of the HH cycle – Na^+^ ions flow into the inner part of the axon, K^+^ channels are still closed. The bottom panel shows the next phase of the HH cycle – Na^+^ channels are already closed, but through now open potassium channels K^+^ ions out-flow to surrounding electrolyte, which causes reduction of the membrane polarization.

In the activation of closed Na^+^ channels various signals compete as stimuli – the opening of these channels is triggered by the fastest one, provided it has the amplitude greater than the required threshold. Other slower stimulus signals cannot initiate the already activated HH cycle because of refractory period, thus only the fastest stimulus can be aided in energy at AP formation, which allows to keep this stimulus amplitude beneath the required threshold over arbitrarily long distance.

The automatic HH cycle, when is ignited by a stimulus, residually strengthens the stimulus and in this way balances its unavoidable Ohmic losses. This must happen at any mechanism of the stimulus (ion diffusive current or plasmon-polariton) in order to assure long range of the AP transduction.

### B. Microscopic structure and functioning of Na^+^ channels by voltage gated

For coupling of plasmon-polariton with HH cycles most important is the triggering of Na^+^ channels by plasmon-polariton stimulus. The latter has high frequency and can open sodium channels only if its frequency is lower than the inverse time scale of triggering these channels. To assess the time-scale of triggering the opening Na^+^ trans-membrane channels, the molecular structure of these channels needs to be taken into account. In recent study of bacteria model of Na^+^ channels [24], the progress in insight into their structure and functioning has been achieved.

The channel for Na^+^ ions gated by a voltage is the complex of polypeptide of ca. 2000 amino acid molecules forming a pore by four almost identical domains located around the pore and mutually connected by intracellular linkers. Each of these domains is composed of six polypeptide helices closely bound by connectors and together passing the cellular membrane – these helices are named as S1 – S6 and are visualized in Figs 6 and 7. Helices S1 – S4 in four modules are tightly located and can deform in response to voltage arisen due to membrane polarization change, whereas the other two helices S5 and S6 take part in construction of the pore (together four such pairs are immediate construction blocks of the quasi-circular pore, cf. Figs 6 and 7). All helices are mutually connected, and S1 – S4 segment is connected with S5 – S6 segment by short linker – small helical spring connecting S4 and S5 (this linker is marked in blue color in Figs 6 and 7). The most sensitive to the vertical trans-membrane voltage is S4 helix (marked in magenta color), which has positive charges locally non-balanced. At the membrane depolarization an electrical force is applied to this charges, which causes a vertical shift of S4 helix (in transverse direction with respect to the membrane and oriented upward or downward depending on the voltage change sign). At steady polarization of the membrane − 70 mV the helix S4 is in its lower position. When the polarization of the membrane changes by ca. 15 mV to − 55 mV, helix S4 quickly moves up a vertical distance of 11.5 Å. This upper position of S4 can be named as its excited state. Due to tight connections between all helices, the shift of S4 causes the conformational change of the whole complex resulting in opening of the channel pore, i.e., in widening of the central collar ring. Such a scenario of opening of the sodium channel [24] supports the earlier sliding helix model [41] associated with translocation of gating charges in the S1 – S4 complex.

**FIG. 6.**
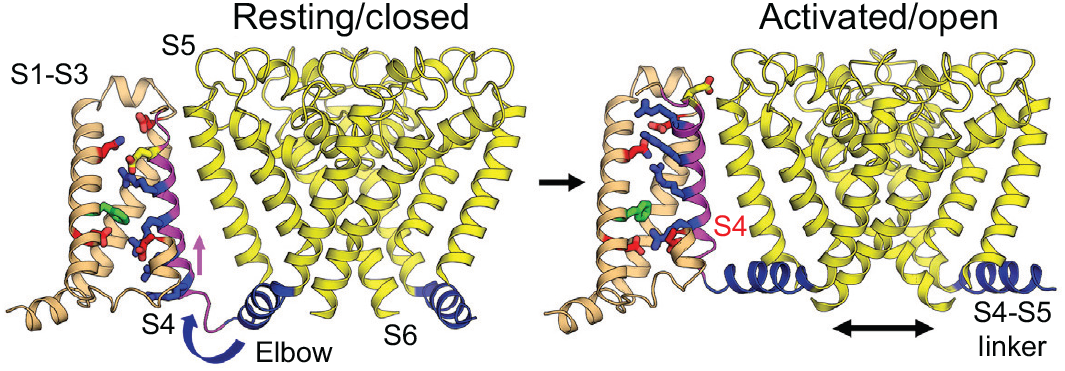
Conformation of polypeptide domain in the sodium channel by voltage gated. The channel is the trans-membrane complex of four domains connected by linkers [24]. The shift and rotation of the helix S4, indicated in magenta color, almost in vertical direction causes opening the tunnel for sodium ions (picture after [24] with permission). The time scale of this conformational change related with the vertical shift of positive charge in the re-polarized membrane when the opening of the channel completes is of order 1 *μ*s [39, 40]. The very precise deciphering of this trans-membrane conformation by [24] agrees with the formerly assumed model for twist and shift of helix S4 [41].

**FIG. 7.**
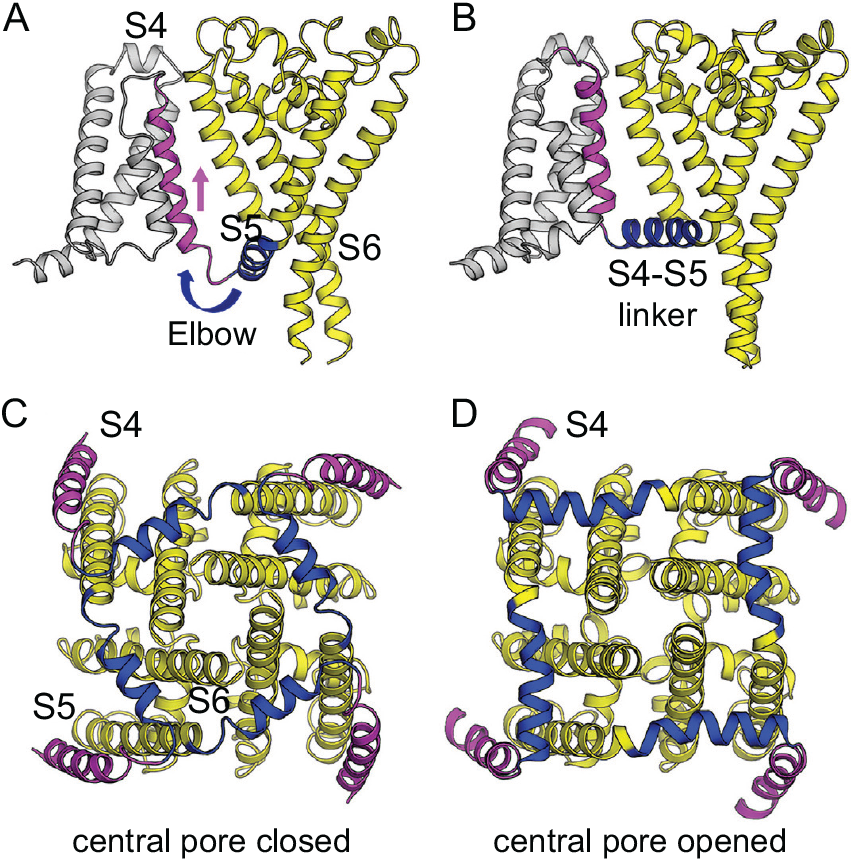
A. Side view of the structure of single complex S1 – S6 (one of the four builders of the sodium channel, as shown in C. and D. in transverse bottom view) focusing on S4 segment (magenta) and the S4-S5 linker (blue), with the S1 – S3 segments shown in gray and the pore module S5 – S6 in yellow, in metastable state at −70 mV membrane polarization. B. In response of depolarization of the membrane (by approximately 15 mV) the S4 segment shifts outward on the span of 11.5 Å and achieves its excited state. The S1 – S3 segments remain unchanged. The S4-S5 linker acts as an elbow that connects the S4 movement to modulate the pore. This linker changes its position and orientation strongly and pulls S5 – S6 segments resulting in opening of the pore. C. Bottom (intracellular) view on the Na^+^ channel with four complexes S1 – S6 (S1 – S3 omitted for clarity) – the state with closed pore in the center. The metastable conformation of the structure at polarization −70 mV is presented – S4 segments are in their unstable lower positions (in magenta color), elbows and linkers S4-S5 are in tension (in blue color), which causes tightening the collar of the pore created by S5 – S6 pair segments. D. Bottom (intracellular) view on the Na^+^ channel with four complexes S1 – S6 (S1 – S3 omitted for clarity) – the state with open pore (central free space with diameter 10.5 Å). The conformation change of the structure is visible – S4 segments shifted upward (in magenta color) pull via elbows the linkers S4-S5 (in blue color), which causes a step type deformation of S5 – S6 pairs and opening of the channel. (Cartoon pictures after [24] with permission).

**FIG. 8.**
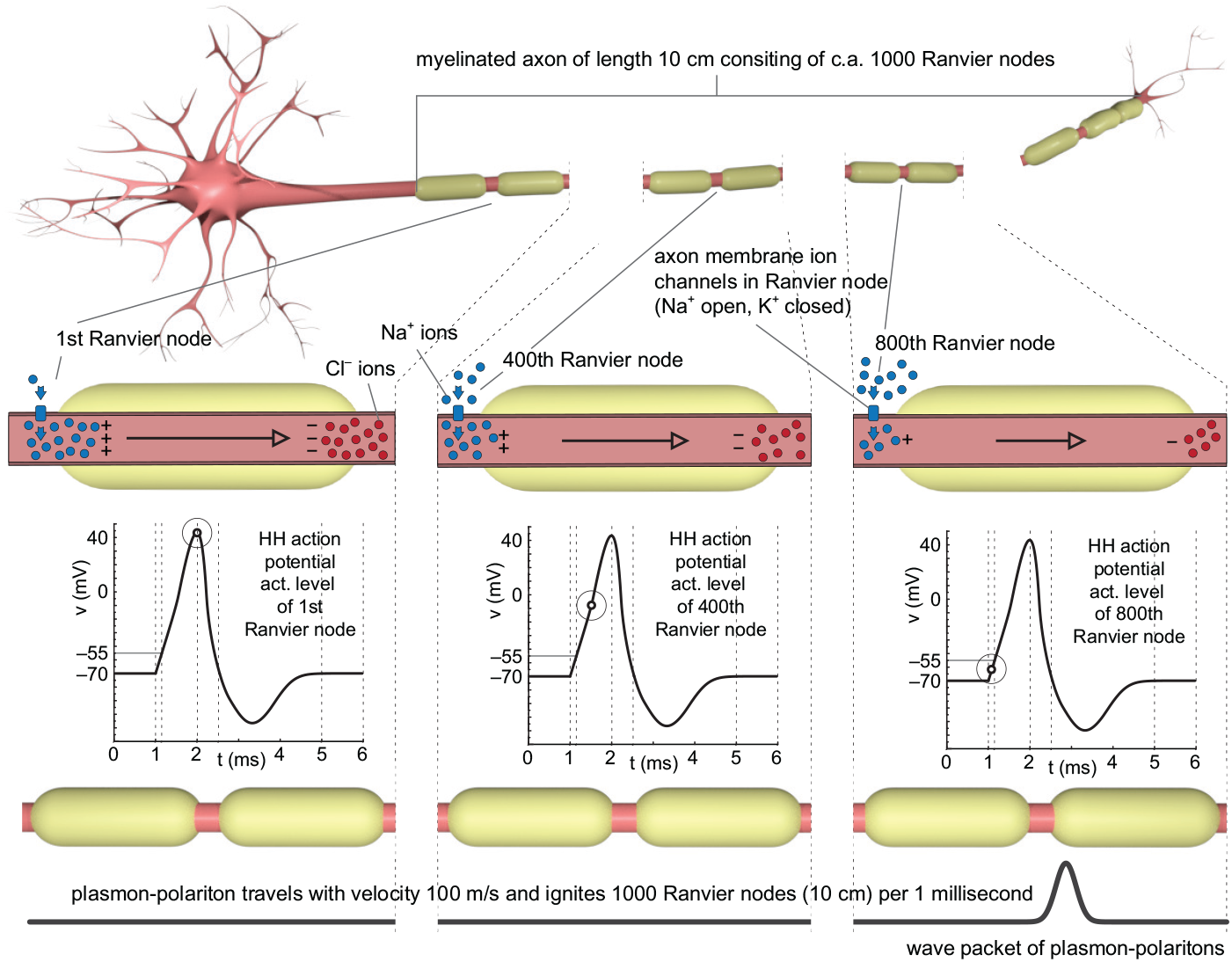
The myelinated axon is a chain of segments, in which can propagate ionic plasmon-polariton – the wave-type synchronized collective oscillations of ion density in segments. This makes a kinetics of the ignition electrical signal without any net diffusion current of ions along the axon. The longitudinal oscillations of electrical dipole on a particular segment of the axon below the myelin sheath is coupled to similar dipole oscillations on other segments in the chain, which, due to periodicity of the whole structure, results in the wave-type signal propagating along the axon with high speed. This wave-type signal is called plasmon-polariton. The wave packet of the plasmon-polariton (cf. III C) triggers the HH cycle on consecutive nodes of Ranvier, so the whole axon fires with the speed as observed at the saltatory conduction. The time retardation of AP creation phase (the phase of the HH cycle) on consecutive nodes of Ranvier is caused by the speed of the plasmon-polariton stimulus signal and temporal scale of HH cycle. Within the time period of 1 ms the stimulus signal triggers 1000 consecutive nodes of Ranvier (as the plasmon polariton wave packet (lower panel) runs with the speed of 100 m/s). For example, the first node is at middle of the HH cycle (the red arrow in left inset), the 400-th node is at early phase of the HH cycle (the red arrow in the middle inset), while the 800-th node just ends its triggering by the stimulus (the red arrow in the right inset).

The molecular system described above is strongly taut by large electrical field forces and tension in helices fixed by hydrophobic forces in several nodes at the rest polarization, − 70 mV. At the steady conditions the whole complex is in a metastable state (unstable local equilibrium) and can be rapidly released in the response to stimulus signal with amplitude exceeding some needed threshold of depolarization of the membrane (the stimulus must change the membrane polarization from − 70 mV to ca. − 55 mV). If the stimulus is an oscillating signal like of plasmon-polariton wave packet, then the period of these oscillations cannot be smaller than the time scale of the rapid shift of S4 helix.

The shift of the S4 helix can be very fast, as we will evaluate below, but some inertia is revealed due to the interconnections between the helices, in particular due to the connection of the S1 – S4 subcomplex to S5 and S6 helices via the linker between S4 and S5. The deformations and shifts of these linkers in four components of the entire channel trigger the jump in conformation of the central part of the channel formed by four pairs of S5 and S6 helices. At the membrane polarization of − 70 mV and lower position of S4, the linkers S4-S5 tighten the collar around all S5 and S6 segments (Fig. 7 C), which prevents opening of the pore formed by four S6 helices (Fig. 7 A). In response to the membrane depolarization by 15 meV, linkers S4-S5 are pulled by the upward shift of S4 helix, which triggers pore opening (Fig. 7 B, D) by the loosening of the collar around centrally located S5 – S6 helices (Fig. 7 D). The rapid shift of S4 segment (magenta in figures) pulls the elbow of the S4-S5 linker (blue). The time scale of this shift and related pore opening is much lower than 1 ms (is rather of the time scale of microsecond). The open pore has the diameter of 10.5 Å sufficiently large to be passed by Na^+^ ions. Influx of sodium ions lasts next ca. 0.5 ms.

When the helix S4 is in the lower position (the spring is tensioned), the collar formed by S4-S5 linkers is tightened and the channel is closed. The lower position of the S4 segment causes the tension which bends the junction between the S4 segment and the linker S4-S5 to the form of inward projecting elbow, which eventually tightens the collar of S4-S5 linkers and narrows the pore to its closed state. This conformation is stabilized by hydrophobic forces in several nodes along the whole structure [24]. This steady-state resembles a cocked gun [24], and a small depolarization causes a quick shot outward of the segment S4. By connection with the S4-S5 linker, the elbow relief loosens the collar and quickly expands the pore. The energy of taut springs of helices in initial unstable equilibrium accelerates the upward movement of S4 segment in addition to electric force of the stimulus. The movement of the four S4 segments rapidly reduces the bend in four elbows connecting to linkers S4-S5 and pulls linkers which eventually open the channel.

The voltage sensitive S4 segment has the mass of ca. 3 *×* 10^−23^ kg and is exposed at triggering to the voltage 15 mV on the distance 11.5 Å, i.e., to the large local electrical field 1.3 × 10^7^ V/m. This field exerts the force (assuming 4 positive elementary charges active) 8 × 10^−12^ N, which causes an acceleration of the mass 3 × 10^−23^ kg equal to 2.7 × 10^11^ m/s^2^. Rough estimation of the time for free shift on the distance 11.5 Å is thus 10^−10^ s. Additional inertia due to chemical binding making the shift not free, slows down the mechanical response by 3 − 4 orders, which results in the time rate for triggering of the opening of a sodium channel of order of 1 *μ*s. For the model we assumed in the present paper the lower limit of this triggering time 1.5 *μ*s (corresponding to angular frequency of plasmon-polariton *ω* ≃ 4.18 MHz obtained via Eq. (7) for maximal group velocity at assumed *ω*_1_ ≃ 5 MHz).

### C. The wave packet of synchronized ion oscillations in myelinated axons and in C fibers carriers the stimulus for HH cycles

#### 1. Competition between stimuli in axons

In an axon there are possible various carriers of the stimulus. Conventionally it is considered a diffusive ion current, which can propagate along any axon (or dendrite) according to the cable model (cf. Appendix A). The mechanism of such a kinetics of charge carriers consists in consecutive charging and discharging of infinitesimal pieces of the cable (inner conducting cord of an axon isolated from an electrolytic surroundings) being RC small circuits mutually coupled in a series. This results in pushing of ions in longitudinal direction along the axon cord, which satisfies the diffusion equation. The velocity of this diffusion is, however, restricted by values of the trans-membrane resistivity and capacity and the longitudinal conductivity of the inside axon cytosol. The realistic values of these parameters range the velocity of the diffusion current to ca. 5 cm/s in humans (for 1 *μ*m diameter of the axon, and this velocity scales with the axon diameter 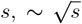, cf. Appendix B), cf. e.g., [9] page 34. Note, that for ultra-thin C fibers with the diameter of order of 0.1 *μ*m, the speed of ion diffusion in the cable-model is thus of order of 1.5 cm/s, far insufficient to clarify the observed micro-saltatory conduction in C fibers with the speed of several meters per second.

In the case of myelinated axons, the thick myelin lipid layer decreases the trans-membrane electrostatic capacity, but increases also the resistivity across the membrane covered by the lipid layer. In the result, one can obtain the increase of the velocity in the cable model by factor 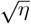, where 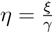 means the ratio of the myelin layer thickness to the thickness of the bare axon membrane. For observable value of *η* ∼ 100 the increase of the signal speed is by factor 10, so that it gives not more than ∼ 1 m/s (for *s* ∼ 1 *μ*m), i.e., still by two orders of magnitude too small in comparison to the observed saltatory conduction speed in such myelinated axons of ∼ 50 ℒ 150 m/s [9–19].

Another mechanism, bereft of slow net ordinary current of ions, can be synchronized oscillations of local ion density in small segments along the axon. Such oscillations can mutually couple in neighboring segments, which results in a very speedy signal along an axon propagating as a wave-type collective synchronized ion oscillations – ion plasmon-polariton. The group velocity of this wave is not limited by the longitudinal resistivity but is strongly controlled by the frequency of charge density oscillations in segments of an axon. The obligatory condition to organize such a stimulus signal transduction along an axon is the periodicity of its longitudinal structure and the synchronization of the frequency of plasmon-polariton with the time-scale of triggering of opening of Na^+^ channels to be capable to ignite HH cycles.

### D. The regulatory role of the thickness of myelin sheath in nervous signaling

It is known from observations, that the thickness of the myelin layer is the major determinant of the proper functioning of myelinated axons. This thickness is greater than that needed for only electrical isolation because the role of myelin is different, not related with isolation only. Though a thicker myelin can reduce the capacity across the membrane covered by myelin, this is, however, insufficient to accelerate the stimulus to the required level, despite a common belief upon the cable model. In the case of plasmon-polariton the role of myelin is different.

The oscillations of ions in myelinated segments interact with surrounding intercellular electrolyte, i.e., the dipole oscillating in the axon cord fragment excites the inversely directed also oscillating dipole of ions in the outer electrolyte – in the electrolytic cave surrounding the myelinated segment, cf. Fig. 9. The coupling of both these dipoles across the myelin sheath with the thickness *ξ*, is of the near-field type because of the closeness of both interacting dipoles [34, 35]. If by *d*_1_(*t*) and *d*_2_(*t*) we denote the longitudinal dipoles in the axon rod in this myelinated segment and in the corresponding cave of surrounding electrolyte, respectively (cf. Fig. 9), then the equation for the dynamics of this coupled system is as follows,

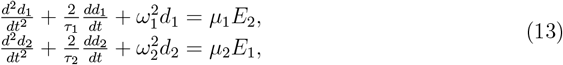

where, 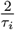 is the damping rate of *i*-th dipole (*i* = 1, 2), *ω*_*i*_ is the self-frequency of each dipole if are separated, *μ*_*i*_ is the longitudinal polarizability of the axon rod segment for *i* = 1 and of the cave in surrounding cytosol for *i* = 2 (the polarizability for a sphere equals to 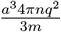, *a* is the sphere radius, *n* is the concentration of ions, *m* is the ion mass, *q* is the ion charge, but for an elongate geometry is anisotropic and different in the longitudinal and transverse direction; we consider here the longitudinal polarization of dipoles). Both dipoles mutually interact and together they form a pair of coupled oscillators. Dipoles couple via their electromagnetic field radiation according to Eq. (1) with only 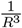terms maintained, corresponding to the most important near-field zone at small distances. The electrical fields induced by dipoles in the near-field zone are thus as follows,

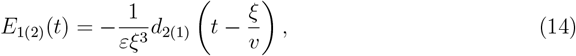

where *ξ* is the thickness of the myelin sheath, 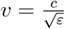 is the light velocity in the electrolyte, *ε* is the dielectric permittivity of the electrolyte, indices 1 and 2 refer to dipoles 1 and 2, respectively. The time retardation on small distances *ξ* is negligible here, thus 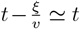. The fields *E*_1(2)_ allow to determine the mutual interaction of dipoles *d*_1_ and *d*_2_ in the near-field coupling approximation if are multiplied by longitudinal polarizability of the rod segment and the cave, respectively (as written in Eq. (13) in the right-hand side).

**FIG. 9.**
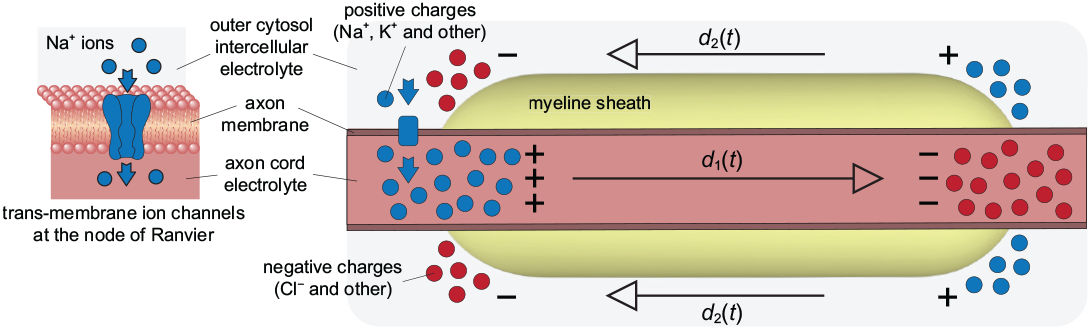
Cartoon of a single myelinated segment – polarization of longitudinal electrical dipole of the inner cytoplasm inside the axon segment under the myelin. To the right oriented arrow, the dipole *d*_1_(*t*) induces local opposite polarization of the outer cytoplasm cave around the myelin segment – to the left oriented arrows of the opposite dipoles, which sum up to the dipole *d*_2_(*t*) also along the axon axis due to axial symmetry. Both dipoles *d*_1_ and *d*_2_ oscillate as the pair of oscillators coupled across the insulating myelin layer. The coupling is weak for sufficiently thick myelin sheath and causes the slow beats due to oscillator coupling. These slow self-oscillations of coupled dipoles lead to a similarly slow frequency of plasmon-polariton wave packet, which can be synchronized with the time scale of the triggering the opening of sodium channels. The thinning of the myelin sheath increases beating frequency and precludes the synchronization with opening-time-scale of sodium channels. The thickening of the myelin sheath is also inconvenient, because for the frequency of beating lower than the optimal one, the plasmon-polariton slows down (as shown in Fig. 3). The optimal thickness of the myelin sheath is thus selected by the synchronization of the plasmon-polariton frequency with the time scale of the triggering the opening of sodium channels.

To analyze the most important feature of the solution of Eq. (13) let us simplify it by the assumption, *μ*_1_ = *μ*_2_ = *μ, ω*_1_ = *ω*_2_ = *ω*_0_, and neglect the damping and the time retardation. The simplified Eq. (13) attains the form,

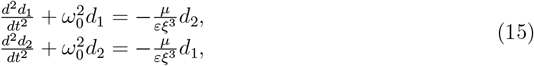

This is the equation for two coupled harmonic oscillators in this simplified case without damping. It is the system of linear differential equations, which can be easily solved in Fourier domain, i.e., assuming the solution in the form, *d*_1_(*t*) = *Ae*^*i*Ω*t*+*ϕ*^ and *d*_2_(*t*) = *Be*^*i*Ω*t*+*ψ*^. The self-frequencies of the coupled system (15) are the nodes of the characteristic determinant of the system (15),

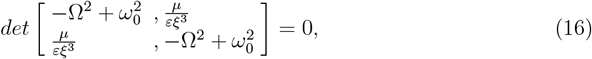

i.e., 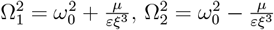.

For the initial condition suitable for the excitation of the considered segment of the axon, i.e., *d*_1_ (0) = *d*, *d* (0) = 0, 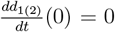, the solution of Eq. (15) is the linear combination of both self-frequency oscillations,

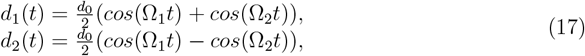

or, equivalently,

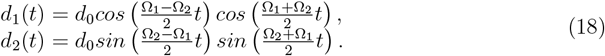

We see that we deal here with the beating with low frequency,

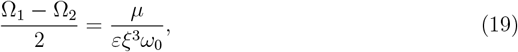

in this notation *ω*_0_ = *ω*_1_ for self-oscillations of myelinated segment if surrounding is neglected. For sufficiently large *ξ* we get thus slow oscillations required for the time scale of the triggering of opening of Na^+^ ion channels at nodes of Ranvier to initiate the HH cycle (cf. Fig. 10). The period of the stimulus signal (with an amplitude beyond the threshold for Na^+^ ion channel opening) cannot be lower than the characteristic time for the triggering the activation of Na^+^ channels. As the beating frequency 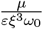 is also inversely proportional to *ω*_0_ (*ω*_0_ = *ω*_1_ here), thus for shorter myelinated segments with higher *ω*_1_, thinner myelin layer gives the same 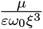, i.e., lower thickness of the myelin assures the same frequency sufficient to trigger the sodium gates at nodes of Ranvier for shorter myelinated segments, which agrees with observations.

**FIG. 10.**
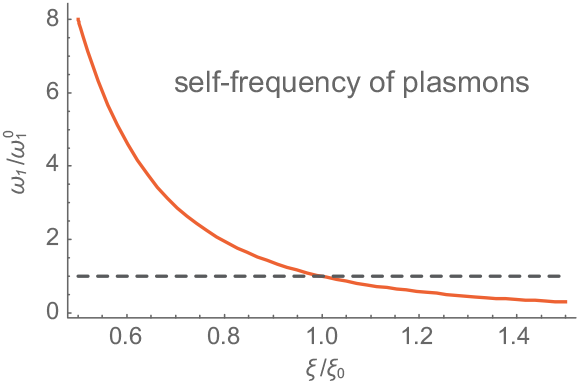
Dependence of the self-frequency of plasmon oscillations in a single myelinated segment, *ω*_1_, versus thickness of the myelin sheath *ξ*, where 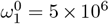 Hz corresponds to the thickness *ξ*_0_ (at which *v*_*z*_ = 140 m/s). A sharp increase of the oscillation frequency is visible with diminishing of the thickness *ξ* (the diminishing by only 20% enhances the frequency twice, by 50% thickness reduction causes 8-fold increase of the frequency).

The frequency of longitudinal plasmon is reduced in a strongly elongated axon rod segment under myelin with the aspect ratio ∼ 0.01 from bulk ion plasmon frequency at ca. 100 mM concentration of ions, 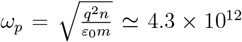 1/s by the factor ∼ 10^2^ and additionally is reduced by an appropriate increase of *ξ* due to the beating effect and finally

achieves the value of ∼ 5 × 10^6^ 1/s, resulting in the saltatory conduction velocity ∼ 150 m/s (as visible in Fig. 3). Thus we see, that the myelin layer thickness controls the velocity of the saltatory conduction via the accommodation of the frequency of the stimulus signal (of plasmon-polariton wave packet) to the time scale of the triggering of opening of Na^+^ channels at nodes of Ranvier (which is of order of a microsecond, cf. paragraph IV B). The synchronized frequency is selected and the corresponding plasmon-polariton wave packet is strengthened and narrowed by the side effect of the HH cycles of AP formation, which repeatedly adds the polarization impulse on consecutive nodes of Ranvier. Such a repeating stimulation of plasmon-polariton modes narrows the wave packet of the plasmon-polariton centered at the optimal mode (the more uniform distribution of the excitations along the chain causes the more narrow shape of the envelope in the wave packet, which is the manifestation of the Fourier representation of a constant function in the form of the Dirac delta). This is in contrast to other non-synchronized frequencies of plasmon-polariton modes. Non-synchronized modes are quickly damped due to Ohmic losses at ion scattering because they are not supported in energy.

The described mechanism of the selection of the sufficiently low self-frequency of plasmon oscillations of the myelinated single segment allows for the precise accommodation of the frequency of plasmon-polariton. Not all frequency modes of plasmon-polariton are persistent. Ohmic damping causes their attenuation on short distances unless the energy is supplied as the by-product at repeating AP formation at consecutive nodes of Ranvier. This requires, however, a perfect coincidence of the plasmon-polariton mode frequency with timing of triggering the opening of Na^+^ channels across the axon membrane at nodes of Ranvier. Too quick oscillations are not able to trigger the opening of sodium channels, which eliminates corresponding modes. Too slow oscillations reduce the velocity of corresponding plasmon-polariton modes and are retarded with respect to the optimal faster one and are late on nodes of Ranvier already ignited by the fastest one. This simple mechanism selects the optimal and persistent wave packet of plasmon-polariton with the velocity observed at the saltatory conduction. This wave packet is supplied with the energy as the by-product at the automatic AP formation at consecutive nodes of Ranvier (eventually on the cost of ATP/ADP cellular mechanism at the HH cycles).

The thickness of the myelin sheath is a control factor for synchronization of plasmon-polariton with the AP formation mechanism needed to balance the Ohmic losses of the plasmon-polariton stimulus signal over arbitrary large distances and allowing for firing the whole axon of arbitrary length with high speed of the saltatory conduction (i.e., firing of ca. 1000 nodes of Ranvier per 1 ms at 100 m/s saltatory conduction speed). Reducing the myelin sheath thickness increases the frequency of the plasmon-polariton which causes its desynchronization with HH cycles and perturbs plasmon-polariton kinetics via its damping, just like in multiple sclerosis.

## V. PLASMON-POLARITON WAVE PACKET WITH MEGAHERTZ FREQUENCY

### A. Frequency of plasmon polariton modes for plasmon frequency 5 MHz in single segment of axon if separated

For plasmon self-frequency in each segment (if separated), *ω*_1_ ≃ 5 × 10^6^ 1/s (5 MHz), one can determine the frequency band *Reω*_*k*_ (for short written next as *ω*_*k*_) for plasmon-polaritons modes for *k* ∈ [0, 2*π/d*) (via the solution of Eq. (7)) as shown in Fig. 3 left panels for a choice of segments length *L* = 100 *μ*m, with aspect ratio 0.01 and for exemplary segment separation *δ/L* = 0.005, 0.05, and 0.1 (corresponding to Ranvier node lengths *δ* of 0.5, 5, and 10 *μ*m, respectively). The derivative of the obtained frequencies *ω*_*k*_ with respect to *k* determines the group velocity of the plasmon-polariton modes. This velocity also depends on *k* and the maximal its value (for appropriately selected wave packets) achieves 100 − 150 m/s, required to explain the saltatory conduction. The results are presented in Fig. 3 (right panels). The maximal value of the group velocity gives *k* and corresponding frequency of collective plasmon-polariton *ω*_*k*_ ≃ 4 MHz (for *ω*_1_ = 5 MHz). Note that such a frequency of ion oscillations in myelinated axon has been actually observed [22], which strongly supports the model. To the frequency of 4 MHz corresponds the time scale 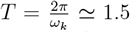 *μ*s. This time scale agrees with minimal time duration of stimulus needed to trigger the opening of Na^+^ channels (as has been demonstrated in Appendix IV B). Thus, the fastest plasmon-polariton mode has the frequency ca. 4 MHz found at *ω*_1_ = 5 MHz frequency of plasmons (i.e., of ion density oscillations) in a single segment (if considered as separated – the mutual coupling between segments in collective plasmon-polariton reduces its frequency to ca. 4 MHz). Higher frequency plasmon-polaritons (though have higher group velocity) cannot open Na^+^ channels and are not supplied in energy by HH cycles at nodes of Ranvier, thus are eliminated by damping on short distances. Lower frequency plasmon-polaritons (despite can open Na^+^ channels) are slower than the optimal one and are insignificant, because Na^+^ channels are opened by the optimal faster signal carrying stimulus with the frequency of ca. 4 MHz. Because of refractory period in HH cycle time-retarded signals are ineffective.

Note, that thicker myelin sheath can be used, however, in the case when some slowing of the saltatory conduction is convenient for physiology reasons, to control over signal retardation in some special complexes of neurons.

From exemplary numerical simulation of plasmon-polariton kinetics for various geometry and electrolyte parameters for axons, we have found that the group velocity of the plasmon-polaritons easily reaches 100 − 150 m/s – cf. Table I.

In Table I a variety of examples are presented, from which is visible that the plasmon-polariton stimulus velocity easily reaches 100 − 150 m/s for realistic values of axon parameters.

From Figs 3 and 4 (and from Table I) we see that it is possible to arrange the wave packet (by selection of appropriate subset of wave numbers *k* to the wave packet, cf. paragraph III C), which can propagate with high velocity. The selection of modes contributing to a wave packet depends on the excitation type. For a point-type localized initial excitation (like a post-synaptic impulse or an impulse from the neuron hillock), all modes *k* are equally excited. Repeated in time the episodes of additional strengthening of plasmon-polariton on consecutive nodes of Ranvier (due to HH cycles) narrows the packet around the fastest mode but only if it is synchronized with timing of the triggering of opening of Na^+^ channels. This latter requirement selects frequency of modes and thus *k*. The mechanism which selects the appropriate *k* range contributing to the wave packet is the damping of those components, which are not synchronized with the HH cycles at consecutive nodes of Ranvier.

The frequency of 4 MHz of plasmon-polariton wave packet allows for the synchronization of dipole oscillations in segments with 1.5 *μ*s time-scale of Na^+^ channel triggering and simultaneously defines the highest possible speed of plasmon-polariton able to ignite the whole axon (the speed of plasmon-polariton grows with its frequency, as shown in Figs 3 and 4). To assure this synchronization the thickness of the myelin must be optimal – the smaller thickness of the myelin accelerates the plasmon-polariton frequency (and its speed) precluding, however, the synchronization, the higher thickness decreases the plasmon-polariton speed. Thus, the mode of plasmon-polariton at optimal thickness of the myelin is stabilized in contrast to other modes with larger or lower frequencies.

The central role plays the time scale of the triggering of the opening of Na^+^ channels. This time scale is defined by the molecular structure of these channels as presented in paragraph IV B. Gaining control over the time scale for the triggering of the opening of sodium channels could be thus a new point for a treatment in the case of demyelination syndromes.

#### 1. Fitting the plasmon-polariton kinetics to the myelinated axon parameters

The bulk plasmon frequency (in infinite electrolyte) [21, 42] is 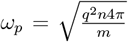 (in Gauss units), or in SI unit system,

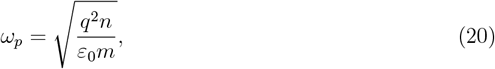

where *q* (in coulombs, C) and *m* (in kg) are the charge and the mass of ions, respectively, *n* is the ion concentration (the number of ions in unit volume, 1 m^3^ of the electrolyte), 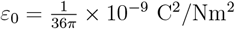 is the dielectric constant. All the estimations will be done here in SI units.

We assumed for the model that the ion charge is *q* = 1.6 × 10^−19^ C and the ion mass is *m* = 10^4^*m*_*e*_, where *m*_*e*_ = 9.1 × 10^−31^ kg is the mass of an electron. For exemplary concentration *n* = 1.9 × 10^16^ 1/m^3^, one obtains the frequency *ω*_*p*_ = 1.8 × 10^7^ 1/s. For ionic dipole surface plasmon oscillations in the spherical geometry of confined finite electrolyte system (the so-called Mie-type frequency for surface dipole plasmons on a sphere, [43]), 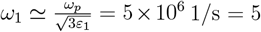 MHz, where the relative permittivity of water is *ε*_1_ ≃ 80 for frequencies in the MHz range, [44] (though for higher frequencies, beginning at approximately 100 GHz, the value of *ε*_1_ of water decreases to approximately 1.7, corresponding to the optical refractive index of water, 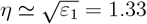). The obtained exemplary *ω*_1_ = 5 × 10^6^ 1/s gives the period *T* = 1.5 *μ*s, as required for the time of triggering of sodium channel opening (cf. paragraph IV B).

However, for the molarity typical for electrolyte in neurons of order of 100 mM (molarity of 1 M = 1 mol/dcm^3^) the density of ions is larger and is of order of 6 × 10^25^ 1/m^3^, which gives *ω*_*p*_ = 4.3 × 10^12^ 1/s and *ω*_1_ = 2.8 × 10^11^ 1/s. These values are too high to accommodate plasmon-polariton frequency (of order of *ω*_1_, more precisely 0.8*ω*_1_ as it follows from the solution of Eq. (7) for *k* corresponding to maximal group velocity, 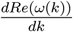) to the time-scale of opening of sodium channels at nodes of Ranvier, for which angular frequency *ω*_1_ must be of order of 5 × 10^6^ 1/s at most (which gives required *T* = 1.5 *μ*s, cf. paragraph IV B). Thus, we must identify paths along which the plasmon-frequency is lowering. If one takes into account the anisotropy of dipole plasmon surface oscillations in highly prolate axon segment with aspect ratio 0.01, then the frequency *ω*_1_ for longitudinal oscillations must be reduced proportionally to the aspect ratio, i.e., to the level of 10^9^ 1/s. Next is the role of the myelin – beating frequency caused by weak coupling of inner cytosol fluctuation with outer electrolyte accros the thick myelin layer can be precisely tuned to the required level of frequency, 10^6^ 1/s, by the appropriately selected myelin thickness. For example, the myelin sheath with typical thickness *ξ* = 0.2 *μ*m (i.e., 0.2 *×* 10^−6^ m) decreases the frequency at the bare cellular membrane (with thickness of *γ* ≃ 7 nm = 7 *×* 10^−9^ m) by the factor 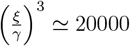, which is sufficient to lower the frequency of plasmon on a single myelinated segment to the scale of 5 MHz.

As the dependence of the beating frequency on the myelin layer thickness is very sharp, 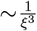 (cf. Fig. 10 and Eq. (19)), this tuning channel is very effective and precise. This elucidates also the fact that even partial deficiency in myelin thickness (as happens at various stadia of Multiple Sclerosis) can severly perturb saltatory conduction, just by the desynchronization of the still fast plasmon-polariton with the HH cycles.

There exists also an additional channel for reducing the frequency of plasmon oscillations in a single segment of an axon. The coupling of plasmon oscillations with electrolytic surroundings causes a non-radiative damping of plasmons, the stronger the thinner the insulating layer is. The energy dissipated into the neuron surroundings is eventually converted into a Joule heat. This effect causes reducing of the *ω*_1_ frequency, according to the ordinary damped oscillator scheme,

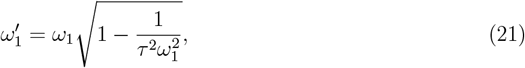

where 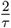 is the overall damping rate for plasmons in a single chain segment including the energy dissipation in the surrounding electrolyte. Similar effect has been analyzed in the case of plasmon-polariton in a metallic nano-chain surrounded by a semiconductor [45]. For myelinated axons this channel of the frequency reduction is negligible due to separation of inner and outer electrolytes by relatively very thick lipid myelin layer, but in the case of naked axons it starts to be important. Especially important is for sufficiently thin axons with dominating surface against volume (the surface of axon segment scales as *s*, whereas its volume as *s*^2^, which favors the surface at low axon diameter *s*). Because the coupling between inner and outer electrolytes undergoes via the separating surface, thus is stronger for lower *s* – such an effect will be discussed in the next paragraph V B with regard to micro-saltatory conduction in naked, but ultra-thin axons of C fibers of pain sensation, where the reduction of plasmon-polariton frequency to the range of 5 MHz is achieved by damping of plasmons according to formula (21).

### B. Micro-saltatory conduction in C fibers

Recently it has been discovered that in ultra-thin (ca. 0.1 *μ*m for diameter) non-myelinated axons in the peripheral neural system for pain sensation (C fibers), the trans-membrane Na^+^ channels cluster on small lipid rafts periodically distributed along axons, as shown schematically in Fig. 11. This localized concentration of Na^+^ channels resembles in structure the ion channel organization at nodes of Ranvier in thicker myelinated axons, and this translates into an equivalent phenomenon of saltatory conduction or related-functional benefits and efficiencies (called in C fibers the micro-saltatory conduction). Observations indicate that AP signal transduction velocity in C fibers by three orders of magnitude exceeds the theoretical cable model estimations is so-thin axons [20].

**FIG. 11.**
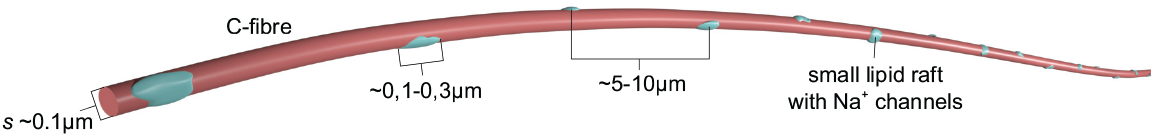
Schematic presentation of C fiber with periodically distributed small lipid rafts with trans-membrane Na^+^ channels.

C fibers are very thin and long non-myelinated peripheral axons responsible for transmitting nociceptive pain sensations (cf. references in [20]). The periodic clustering of Na^+^ channels on lipid rafts resembling the nodes of Ranvier in myelinated fibers are typically of length 0.1 − 0.3 *μ*m and span 5 − 10 *μ*m. They are suggested to permit micro-saltatory conduction in those thin axons despite them being non-myelinated [20]. This emphasizes that the central role for the saltatory conduction plays the periodicity imposed on the linear structure of a net axon. The periodicity allows the wave-type propagation of a stimulus signal in the form of ion plasmon-polariton wave packet which ignites the consecutive blocks of Na^+^ channels where the formation of AP takes place according to HH mechanism (K^+^ channels are still randomly distributed along C fibers). The synchronized ion oscillations in periodic internodal segments (between consecutive lipid rafts along the C fiber) can be organized without any myelin sheath. The myelin plays only the regulatory role as described above to precisely accommodate the plasmon-polariton mode frequency to the timing of the triggering opening of Na^+^ channels.

The AP formation at lipid rafts (similarly as at nodes of Ranvier) is of major importance for the selection of an appropriate mode of plasmon-polariton – that one which is synchronized with HH cycles at consecutive nodes. Only synchronized modes with HH cycles will be persistent due to energy supplementation by the occasion of AP formation on consecutive rafts. Other, non-synchronized modes are quickly eliminated due to strong damping of plasmon-polaritons on short distances, as not supplemented with energy. For the triggering of opening of N^+^ channels the frequency of plasmon-polariton stimulus cannot be larger than the inverse of the lowest time span needed to trigger the opening of Na^+^ channel. This condition is satisfied by ca. 4 MHz frequency, thus higher frequency modes cannot be synchronized with HH cycles and without energy supplementation are eliminated bereft any communication effect. Lower frequencies can be synchronized with the triggering the sodium channel activity, but for them the group velocity of plasmon-polariton mode is smaller, so they are retarded in comparison to the fastest one and also do not ignite HH cycles at consecutive nodes (rafts), because these cycles are already ignited by the fastest synchronized mode.

This scenario explains the micro-saltatory conduction in C fibers. The absence of the myelin sheath causes, however, that the frequency of the plasmon-polariton in C fibers would be larger than in myelinated axons. Instead of myelin the frequency of plasmon-polariton is mitigated in C fibers by the strong damping of ion oscillations in periodic segments. This damping is caused by the Ohmic losses large in the inner neuron cytosol with small diameter *s* and by the strong coupling to outer electrolyte across the naked membrane. The latter is especially large at the absence of a myelin as in C fibers and for very thin axons, for which the role of the surface versus volume of the axon is enhanced, thus favors the surface effect – the electric coupling with surrounding electrolyte across the membrane. Thus, for these thin and naked fibers we get the self-frequency for plasmons at a single segment according to Eq. (21), where 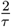 represents the overall damping rate (Eq. (21) displays the frequency of an ordinary harmonic damped oscillator). For large damping the reduction of the frequency is very efficient (this favors, however, ultra-thin bare axons), but the frequency 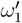 still can be larger than required for the synchronization with the Na^+^ channel triggering time. In such a situation it will be selected the mode *k* of plasmon-polariton with as much as possible lowered frequency of plasmon-polariton as in Fig. 3 (left panel, left low corner). This gives not the extreme speed for the micro-saltatory conduction acc. to Fig. 3 (right panel), though highly exceeding the diffusion of ions which is particularly small in ultra-thin axons bereft of myelin. This agrees with observations of the speed of micro-saltatory conduction in C fibers of order of several meters per second, whereas the diffusion current here has the velocity of 1.5 cm/s, at most. Note that the invention of the myelin reduces transmembrane energy losses and, on the other hand, gives the mechanism of lowering of plasmon-polariton frequency precisely controlled by the myelin thickness in myelinated axons, quite conversely to C fibers. In myelinated axons the maximal velocity mode (the maximal in Fig. 3, right panel) can be synchronized with Na^+^ channels and HH cycles. Therefore, in the case of C fibers the micro-saltatory conduction does not reach as high speed as in the myelinated axons, but still by three orders of magnitude larger than the diffusion current (very slow in ultra-thin axons).

It is thus evident that the cable theory (coupled with HH cycle of AP formation) is helpful only in short dendrites and unmyelinated axons without any periodic structure of Na^+^ channel distribution. The periodicity (by the myelination separated by nodes of Ranvier or by periodic rafts with Na^+^ channel clusters) always causes plasmon-polariton kinetics, which if synchronized with HH cycles, is much more efficient and energy frugal, persistent and fast over arbitrary long distances. We note, however, that no problem is the high velocity of a plasmon-polariton signal, which can be very large for high frequency modes. The problem is that such high frequency modes cannot trigger the opening of Na^+^ channels and cannot be synchronized with HH cycles, thus are quickly eliminated as damped without energy supplementation. Only synchronized with HH cycles plasmon-polariton modes can propagate on long distances despite strong damping because of the energy supplementation at each node with HH cycle of AP formation.

## VI. DISCUSSION AND CONCLUSIONS

The developed in the present paper plasmon-polariton model of the saltatory conduction in myelinated axons and in C fibers of pain sensation seems to pave a new way for better insight into neuro-signaling with some consequences for the potential improvement of future treatment of demyelination syndromes and for fighting pain. The satisfactory agreement of the plasmonic model with observations has been achieved for actual neuron parameters, which was inaccessible for the conventional previous approach upon the cable model. Plasmon-polariton model agrees with observations for stimulus signal velocity at saltatory conduction for both myelinated axons and C fibers, and for other properties of this kinetics, like size and temperature scaling and molecular mechanism of Na^+^ channel triggering. The different role for myelin thickness has been identified than that within cable model and the mechanism for control over plasmon-polariton in nonmyelinated C fibers has been also provided elucidating why these axons must be extremely thin.

The presented model of ion plasmon-polariton in axons was formulated with the mathematical rigor in an analytical version enabling its direct verification, easy numerical simulations and further development. The agreement with wide class of observations has been demonstrated and some additional supporting data are indicated, ultimately confirming the model of wave-type plasmon-polariton kinetics for stimulus at the saltatory conduction.

The unique properties of plasmon-polaritons that distinguish these signals from ordinary currents may allow for a better understanding of the function and role of myelinated axons in the central nervous system, which can be additionally commented. Despite such axons constitute majority of axons in the brain and spinal cord, the actual role of the myelin in white matter in brain and spinal cord is still unclear in the context of nonmyelinated axons and dendrites dominating in gray matter of the cortex. As the cerebral cortex is considered as the medium for the mind organization, the information processing in gray matter probably takes advantage from the absence of myelin there. Thus, arises a question, why the white matter is myelinted. The answer probably lies in by two orders higher velocity of AP transduction in myelinated axons due to speedy plasmon-polariton stimulus. Hence, one can question why not all neuron filaments are myelinated, as it is so convenient in acceleration of signals. The answer might be related to other property of plasmon-polaritons – the absence of their external e-m signature in contrast to ordinary AC electrical currents. This property precludes participation of speedy plasmon-polariton signals in induction-type resonances in multi-loop highly braided neuron filament networks. The latter in cortex with ordinary diffusive currents are suitable to such operation, i.e., to identification of highly braided multi-loop circuits of axons and dendrites mutually connected by synapses via induction resonances with similarly braided filament bunches in other parts of the brain (like hippocampus). The braids of neuron filaments can be carriers of information possible to be operated by low frequency brain waves. This would be the reason why cortex is not myelinated. Plasmon-polaritons cannot participate in induction resonances as do not have e-m signature. The high speed of plasmon-polaritons is, however, convenient for communication functions both in central and peripheral neural systems.

Another notice would be addressed to the immunity of plasmon-polariton kinetics to external electromagnetic perturbations. This property is certainly convenient for signals of fundamental significance for life functions of organism. Plasmon-polaritons are naturally protected against environmental electromagnetic perturbations. It would also motivate the observation that the white matter in the spinal cord is the outer layer compared to the gray matter, unlike in the brain where the gray matter layer is outer. The activity of the gray matter in the spinal cord is related to the most important life functions, therefore it is additionally protected against any environmental disorders by a layer of white matter. In the cerebral cortex, the dominant functions are related to thinking-type activity, which does not seem to require additional isolation from the environment.

## Acknowledgments

The paper has been supported by Polish NCN project P.2018/31/B/ST3/03764.

## Appendix A Short derivation of the cable model

The cable model is in fact the specific case of the long line theory within the modern telecommunication (cf. Fig. 12). The long line consists of two components – the inner conductor (a wire) and outer conductor separated by an insulator (frequently in the form of a tube, like in coaxial cables). In general, such a line can be represented by a series of infinitely short segments of length *dz*, as shown in the scheme in Fig. 12. *R* is the longitudinal resistance of the inner line per its length unit, *L* is the distributed inductance of the line per its length unit, *C* is the capacity between two conducting components across the separating dielectric layer also per unit length of the line, *G* is the electrical conduction across the barrier per unit length of the line.

**FIG. 12.**
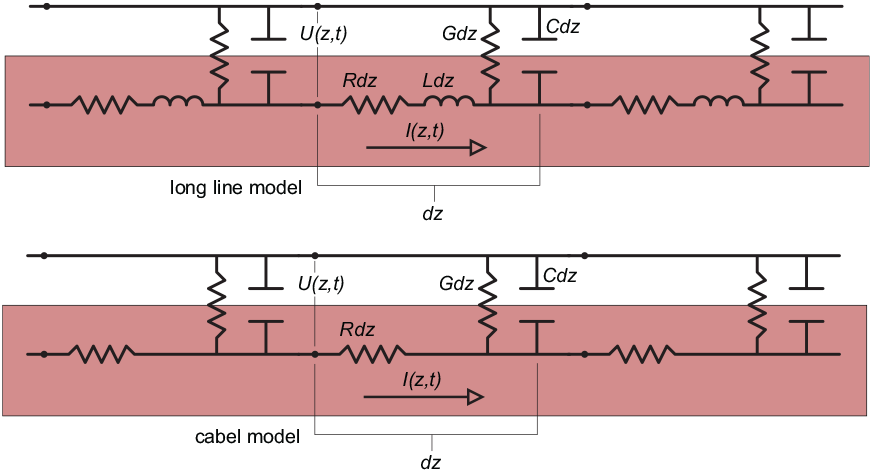
The scheme of the elementary segment with length *dz* for the transmission long line in electro-signaling theory in telecommunication (upper) and the *dz* segment for the cable model (lower). The cable model is a specific case of the long line at the zero inductance.

Due to the Ohm law and the definition of the capacity and the inductance, one gets the relations between the voltage gated the line and the current along the inner wire for a segment with the *dz* length, as in Fig. 12 (upper panel),

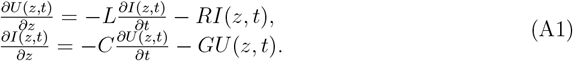

The above equations are two coupled differential equations for the complex voltage *V* and current *I*. Taking the derivative 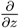 of both equations and substituting one into another, we can arrive in the second order differential equations,

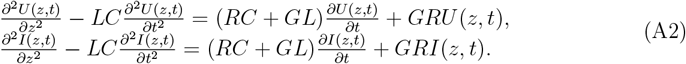

Both equations are identical (they are conventionally called the telegrapher’s equations). For the lossless case, i.e., when *R* = *G* = 0, Eqs (A2) gain the form of the wave equations both for *U* and *I*, describing the ideal wave-type transmission along *z* direction with the velocity 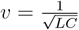 and without any losses. This corresponds to an ideal coaxial line. This is, however, not the case of a neuron, for which the condition *L* = 0 holds (and *R* and *G* are not zero).

The long line for *L* = 0 describes a bare neuron cord – dendrite or axon – separated by a thin insulating cellular membrane from the outer inter-cellular electrolyte. Then, Eqs. (A2) attain the form of 1D diffusion equation both for local voltage *U* (*z, t*) and current *I*(*z, t*),

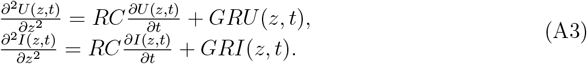

Eqs (A3) constitute the cable model.

Defining the parameters 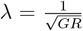 and 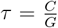, one can rewrite the above equation (A3) (for *U*, the same equation holds for *I*) in a more popular form,

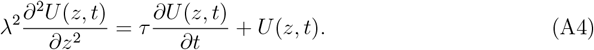

The parameter *λ* defines the spatial scale of the diffusion of ions in the cable, whereas *τ* its time scale. The velocity of the diffusion of the signal, i.e., of the diffusive ion current along the dendrites (or axon) is 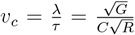. This velocity is the larger the smaller *C* and *R* and the larger *G* are. The range of the diffusion, ∼ *λ*, lowers, however, with growing *G* (due to the shunt escape of the current). Larger *G* results in larger velocity but severely limits the range of ion diffusion.

The overall behavior of the *U* (*z, t*) (or *I*(*z, t*)) diffusion defined by Eq. (A4) can be illustrated as in Fig. 14, which presents the solution of Eq. (A4) for initial condition in the form of Gaussian signal in *z* = 0 point, 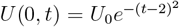. One can notice that the cable theory (i.e., Eq. (A4)) gives the non-wave-type propagation related to an ordinary current (and voltage signal) of pure diffusion type, thus on a relatively short distance with a quickly lowering amplitude (both in position and time domains, what is visible in Fig. 14). For the realistic values of *R, C* and *G* in dendrites or axons, the estimation of *v*_*c*_ gives only few cm/s.

**FIG. 13.**
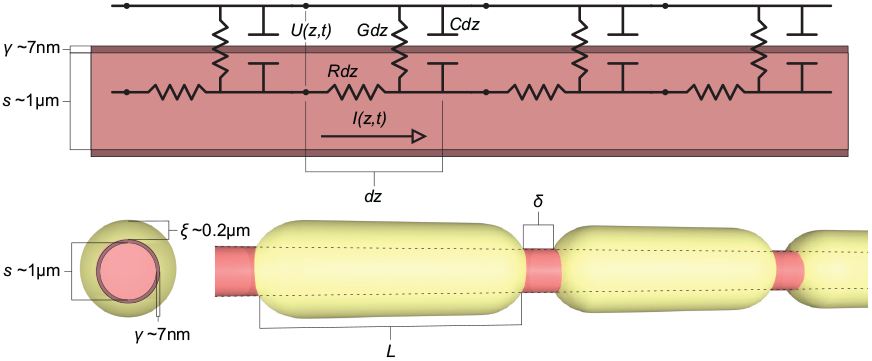
The scheme of the elementary segment with length *dz* for myelinated axon. The thick sheath of insulating myelin decreases trans-membrane capacity *C* and *G* – thus, eventually enhances longitudinal diffusion velocity upon the cable model proportionally to the myelin thickness 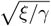.

**FIG. 14.**
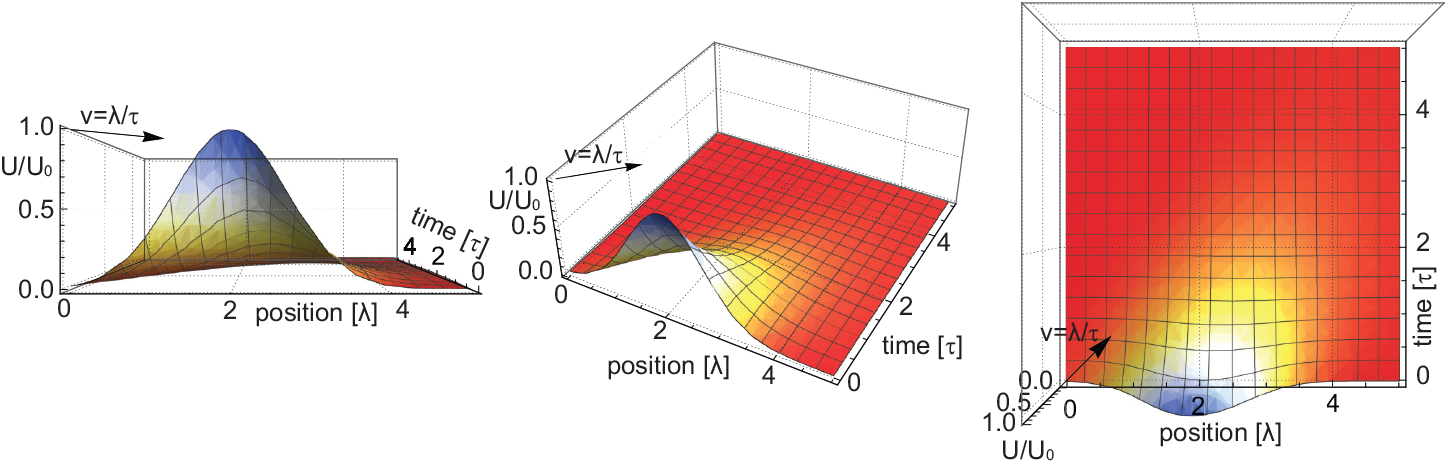
The exemplary solution of Eq. (A4) for diffusion of the ion signal *U* in the cable model for Gaussian ignition at position *z* = 0, 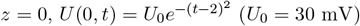 (*U* = 30 mV) and for velocity of the diffusion 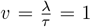 m/s (the graph of *U* (*z, t*) is shown with respect to position *z* in units *λ* and time in units *τ*). The quick vanishing of the signal both in the position and the time domain is visible (on the scale of single *λ* and *τ*, respectively). To assure longer distance of signal conduction the amplitude strengthening is required (at Ranvier nodes), but this does not increase the signal speed. For the saltatory conduction with speed 100 − 150 m/s, in the period of time of 1 ms, 1000 − 1500 consecutive nodes of Ranvier are excited. The majority of them are still in their initial phases in due of this 1 ms, as the AP formation according to the HH model takes time of ca. 1 ms. The diffusive signal with the speed of 1 m/s cannot thus assure the saltatory conduction of the stimulus signal. For realistic axons the cable model signal speed is usually of order at most 1 − 3 m/s (often lower for thinner axons).

The efforts to explain the saltatory conduction in myelined axons upon the cable theory for myelinated axons took advantage of the decrease of the capacity *C* across the membrane covered by a thick lipid myelin layer, cf. Fig. 13. However, this thick lipid layer also causes the decrease of the conductivity *G* across the barrier between inner and outer electrolytes. Both *C* and *G* are inversely proportional to the thickness *ξ* of the myelin layer, thus the speed of diffusive current along the axon, 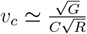 scales as 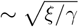. This gives an increase in the velocity of signal by a factor ca. 10 at 100-fold increase of the lipid barrier thickness due to myelination. This is too small to explain the saltatory conduction because myelin reduces the discrepancy from three to two orders of magnitude only.

As the velocity of the diffusive current (upon the cable model) depends also of the diameter *s* of the axon, i.e., *v*_*c*_ scales as 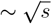 (and even as ∼ *s* in the multi-node HH models [9]), thus, some authors try to increase *v*_*c*_ assuming geometry and size parameters unrealistically large, at least for human cells [12] in contrast to observations [12, 15, 16]. For realistic parameters of myelinated axons the speed of the diffusive current within the cable model is by two orders lower than observed saltatory conduction velocity in myelinated axons [46].

## Appendix B Insufficiency of the cable model to explain the saltatory conduction

For the velocity of a diffusive signal in cable model one gets 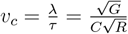, [3, 9–11, 18], as has been derived (cf. Eqs (A3) and (A4) in Appendix A). For the geometry of an axon 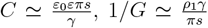 and 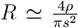, where *s* is diameter of the axon (dendrite), *γ* is the cell membrane thickness (typically, *γ* ≃ 7 − 8 nm), *ρ* is the longitudinal resistivity of the inner cytosol and *ρ*_1_ is the resistivity across the membrane, *ε*_0_ is the dielectric constant and *ε* is the dielectric permittivity of the lipid membrane. Thus, the velocity *v*_*c*_ scales as 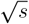. The typical for neuron cells (of humans) the membrane capacity per surface unit, *c*_*m*_ = 1 *μ*F/cm^2^, *ρ*_1_*γ* = 20000 Ωcm^2^, *ρ* = 100 Ωcm, thus, for the diameter of an axon *s* = 1 *μ*m, one gets *v*_*c*_ = 3.5 cm/s [9]. For thinner dendrites *v*_*c*_ is lower proportionally to *s*.

The length of a myelinated segments *L*, the myelin thickness *ξ* and the length of naked breaks *δ* (the length of a node of Ranvier) between consecutive myelinated segments vary in large range and are correlated with the axon diameter (for *s* ≃ 1 *μ*m, typical *ξ* ≃ 0.2 − 0.3 *μ*m, *L* ≃ 100 *μ*m and *δ* ≃ 1 *μ*m). The myelin layer reduces both the capacity and conductance across the membrane covered with this layer, proportionally to the myelin layer thickness.

Hence, the velocity 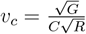, grows e.g., 10 times if the capacitance and conductance lower 100 times (due to 100 times thicker myelin in comparison to the naked membrane). For the above example it gives *v*_*c*_ ≃ 35 cm/s in comparison to ca. 100 m/s observed for the saltatory conduction in myelinated axons of similar size [9–19]. Additionally, the HH cycles at consecutive nodes of Ranvier slow down the speed of the diffusion in the myelinated internodal segments to the level of factor 6 instead of 10 in the above example (at 100-fold increase of the barrier due to the myelin layer thickness) [9]. For realistic (human) axons the numerous simulations of various versions of the cable model interfering with HH cycles on consecutive nodes of Ranvier give at least one order of the magnitude too small speed of signal in myelinated axons in comparison to observations – as illustrated in Table II.

**TABLE II.**
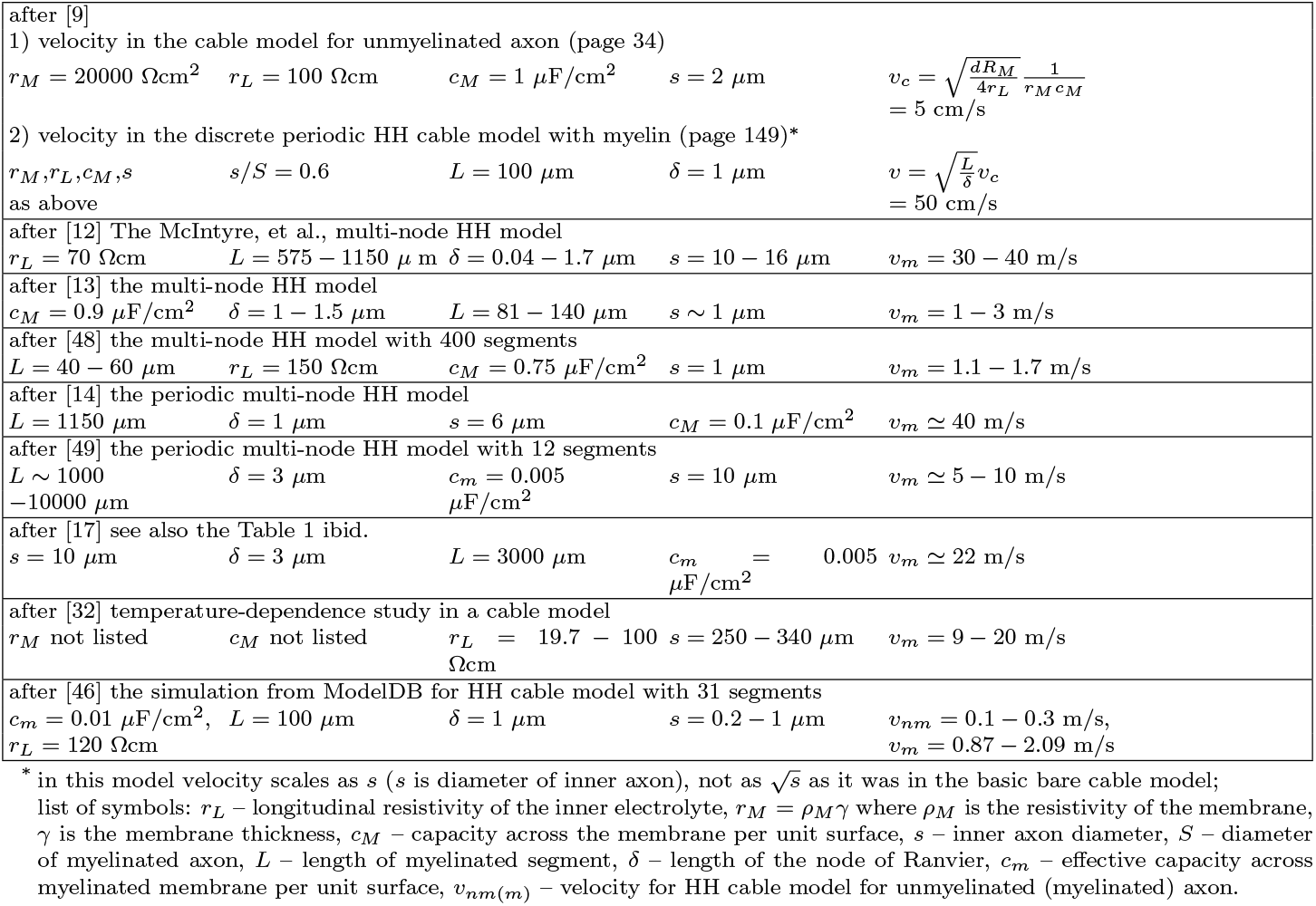
Comparison of simulations within the HH cable model approach.

In the discrete diffusion HH model [9], the velocity in a myelinated axon scales as, 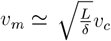, where *L* is the length of internodal segments and *δ* is the length of the node of Ranvier. For instance, in [12, 14] *δ* = 10 *μ*m (and lower) and *L* = 1150 *μ*m (and greater) have been artificially assumed to gain the velocity of the AP transduction, *v*_*m*_ ∼ 40 m/s (cf. also [15–17, 19]), i.e., there are ca. ten times larger in dimension than the actual myelinated axons of a human. It is thus clear that the cable model is not effective in modeling of the observed fast saltatory conduction in myelinated axons. The velocity predicted upon the discrete HH cable model for realistic axon parameters is by at least one order of magnitude smaller than observed [10–19, 47].

The attempts to optimize a model of the cable theory mixed with the HH mechanism in order to obtain a sufficiently high AP velocity in myelinated axons [9–11, 15, 17] did not give a success, which strongly suggests that the way to understand the saltatory conduction must be linked with a different physical mechanism rather than with a lifting of the cable theory of ion diffusion.

The acceleration of signal transmission upon the discrete HH cable model occurs sensitive to various parameters like internodal length of myelinated axons, the size of nodes of Ranvier, myelin thickness, electrical characteristics of neuron membrane and internal cytosol and structure of nodes. It also changes the scaling with respect to axon diameter to a linear one, ∼ *S*, in agreement with observations. Such a behavior was the subject of intensive numerical studies. In Table II nine different simulations of signal kinetics upon the conventional multi-nodal HH cable model are presented.

Symbols for electrical parameters used in Table II slightly differ than previously introduced (to be in compliance with cited references in the Table). If *s* is the diameter of the inner axon, *γ* the thickness of the bare axon membrane, then the longitudinal resistivity per unit length 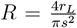, where *r*_*L*_ in Table II is the resistivity of the inner electrolyte (is the same as *ρ* previously used). The capacity across the membrane per unit length, *C* = *c*_*M*_ *πs*, where *c*_*M*_ is the capacitance per unit surface of the membrane. The conductivity across the membrane per unit length, 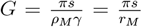, where *ρ*_*M*_ *γ* = *r*_*M*_ (*ρ*_*M*_ is the resistivity of the membrane, previously denoted as *ρ*_1_). Hence, 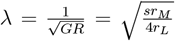 and 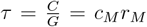. If *r*_*L*_, *r*_*M*_, *c*_*M*_, *s* are given in cgs or mixed units (as in Table II), they must be converted to SI to obtain 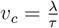 in m/s.

As is visible from the simulations listed in Table II, the velocity of signal in myelinated axons strongly depends on the diameter *s* of an axon (in periodic multi-node HH the velocity is proportional to *s*, whereas in the bare cable model to 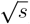) and on the square root of the ratio of internodal length to the node length 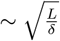. For typical electrical parameters of axon (listed in Table II for each simulation) the realistic velocity within cable model including HH mechanism at nodes of Ranvier does not pass 5 m/s (for human axons being predominantly of order of 1 *μ*m thick [33]), which is, however, by more than one order of the magnitude lower than observed speed of the saltatory conduction in such axons, which varies between 100 m/s to 150 m/s depending of the size of axons [32, 33]. Taking into account that the values for the signal speed of order of 20 − 40 m/s is achieved in periodic discrete HH cable model at unrealistic for humans (and mammals) lengths of the myelinated segments (ca. 1000 − 2000 *μ*m), length of the Ranvier gap (ca. 3 *μ*m) and simultaneously large axon thickness (ca. 10 − 300 *μ*m for the diameter) – cf. Table II, we see, that the cable model even if convolved with the HH mechanism into a discrete multi-node HH model, is unable to fit observations. Note, that the majority of myelinated axons in human central nervous system have diameter lower than 1 *μ*m, [33] and in the peripheral nervous system human axons are thicker (up to 1 − 3 *μ*m in average), but not reach 10 − 20 *μ*m (as assumed in some simulations presented in Table II), which cannot explain the saltatory conduction in humans with a speed of order of 100 − 150 m/s within the discrete cable model.

In the realistic simulation [46] (the last one shown in Table II) for the thin non-myelinated axon with diameter *d* ∼ 0.2 *μ*m, the velocity within the cable model including evenly distributed sodium and potassium channels is ca. 10 cm/s. This is in sharp incompatibility with the observation of so-called micro-saltatory conduction in ultra-thin unmyelinated C fibers of pain sensation in the peripheral nervous system, which is much faster [20]. The increase of the signal speed cannot be associated here with the myelin, as C fibers are non-myelinated.

## Notes

### Competing Interest Statement

The authors have declared no competing interest.

